# Fronto-striatal Functional Connectivity Supports Reward-Enhanced Memory in Older Adults

**DOI:** 10.1101/592501

**Authors:** Holly J. Bowen, Jaclyn H. Ford, Cheryl L. Grady, Julia Spaniol

## Abstract

Both younger and older adults prioritize reward-associated stimuli in memory, but there has been little research on possible age differences in the neural mechanisms mediating this effect. In the current study, we examine neural activation and functional connectivity in healthy younger and older adults to test the hypothesis that older adults would engage prefrontal regions to a greater extent in the service of reward-enhanced memory. While undergoing MRI, target stimuli were presented after high or low-reward cues. The cues indicated the reward value for successfully recognizing the stimulus on a memory test 24-hours later. We replicated prior findings that both older and younger and adults had better memory for high compared to low-reward stimuli. Critically, in older, but not younger adults, this enhanced subsequent memory for high-reward items was supported by greater connectivity between the caudate and bilateral inferior frontal gyrus. The findings add to the growing literature on motivation-cognition interactions in healthy aging, and provide novel evidence of an age-related shift in the neural underpinnings of reward-motivated encoding.

## Frontal-Striatal Functional Connectivity Supports Reward-Enhanced Memory in Older Adults

Reward-associated stimuli receive prioritized attention (Anderson, 2017; Williams et al., 2018, 2017), encoding (Adcock et al., 2006; Castel, 2007; Castel et al., 2002; Cohen et al., 2014; Hennessee et al., 2017; Shigemune et al., 2014, 2010), and consolidation (Spaniol et al., 2014). While there is some evidence that the influence of reward on memory is preserved in healthy aging (e.g., Castel, 2007; Castel et al., 2002; Mather and Schoeke, 2011; Spaniol et al., 2014), there has been little research on age differences in the neural mechanisms mediating reward-enhanced memory. The goal of the current study was to compare reward-enhanced memory in healthy younger and older adults using functional magnetic resonance imaging (fMRI).

Behavioral and fMRI studies have shed light on the influence of reward motivation on declarative memory formation and its neural circuity in younger adults (for reviews, see Miendlarzewska, Bavelier, & Schwartz, 2016; Shohamy & Adcock, 2010). For example, using an incidental memory paradigm, Wittmann and colleagues (2005) found that images serving as reward-predicting cues during a numbers comparison task activated regions of the dopaminergic reward network, including the bilateral putamen, right caudate and bilateral nucleus accumbens, as well as insula and thalamus. Further, higher neural response in midbrain regions to reward-predicting images was linked to successful recognition of these images 3 weeks after encoding. Similar findings have been reported for intentional memory tasks. For example, Adcock and colleagues (Adcock et al., 2006) employed a Monetary Incentive Encoding (MIE) task in which reward was contingent on successful (intentional) encoding of scene stimuli. After a 24-hour delay, high-reward scenes were better remembered than low-reward scenes. This behavioral effect was associated with enhanced connectivity between the hippocampus and regions of the reward network during the presentation of reward cues that preceded the to-be-remembered scenes at encoding.

A small body of work (detailed below) has revealed that healthy older adults also demonstrate reward-enhanced memory (but see Geddes, Mattfeld, Angeles, Keshavan, & Gabrieli, 2018, described in more detail below). Using an incidental memory paradigm, Mather and Schoeke (2011) found that both younger and older adults showed better memory for objects that had been encoded before or after a reward, compared to objects that had been encoded in temporal vicinity to loss or neutral outcomes. Reward also modulates intentional memory formation in older adults. Studies by Castel and colleagues (Castel, 2007; Castel et al., 2002; Cohen et al., 2016; Hennessee et al., 2017) have demonstrated that valuable information is more likely to be remembered than less-valuable information, in both younger and older adults. Finally, in a study using an MIE task similar to that of Adcock and colleagues (2006) we showed reward-enhanced scene memory in both younger and older adults (Spaniol et al., 2014).

The finding of a preserved reward-memory interface in healthy aging is intriguing when considered against the backdrop of age-related decline in the component processes of reward-motivated memory. First, healthy aging is associated with well-documented decline in episodic memory (Cabeza et al., 2018; Grady, 2012; Nilsson, 2003; Nyberg et al., 2012; Shing et al., 2010), so the finding that reward-motivation can boost episodic memory performance in older adults is an important discovery. Second, dopaminergic neuromodulation – thought to play a critical role in reward processing (Arias-Carrión et al., 2010; Berridge, 2007; Berridge and Robinson, 1998; Chowdhury et al., 2013; Ridderinkhof et al., 2012) – undergoes age-related decline (Bäckman, Lindenberger, Li, & Nyberg, 2010; Li, Lindenberger, & Bäckman, 2010). Accordingly, age-related deficits have been reported in stimulus-reward learning (Bäckman et al., 2010; Eppinger et al., 2011), reward-based attention (Williams et al., 2018), and reward-based decision making (Samanez-Larkin et al., 2014, 2011). Interestingly, a recent metaanalysis (Karrer et al., 2017) suggests that although dopamine receptors and transporters are impacted by age, dopamine synthesis appears largely spared in healthy aging. This combination of factors may reduce dopamine re-uptake in older adults, and may partially explain why older adults are able to engage dopaminergic midbrain regions during gain and loss anticipation (Spaniol, Bowen, Wegier, & Grady, 2015; but see Samanez-Larkin et al., 2007, for evidence of reduced loss sensitivity in older adults), as well as in response to gain and loss feedback (Bowen et al., 2019). As Karrer and colleagues (2017) note, it is still not clear, due to a lack of empirical work, why age-related changes in the dopamine system appear to have more impact in some cognitive domains than in others.

Most extant fMRI studies on aging and reward-motivated cognition have focused on characterizing activity within the reward network, with relatively little attention paid to reward-based modulation of other brain regions. In one study, whole-brain analyses of fMRI data acquired during an incentive processing task revealed that older adults, but not younger adults, recruited the lateral prefrontal cortex during processing of high-value gain and loss cues (Spaniol et al., 2015). The extent of lateral frontal recruitment in older adults was associated with higher response speed, possibly reflecting the engagement of cognitive control mechanisms. Age-related over-recruitment of frontal regions is commonly reported in the fMRI literature, and in some cases may serve to compensate for age-related decline in tasks that require high levels of cognitive control (Cabeza et al., 2018; Grady, 2012). However, few published studies to date have examined age differences in reward-based modulation of tasks with high control demands. In particular, there have only been two studies looking at fMRI correlates of reward-modulated long-term memory encoding in younger and older adults (Cohen et al., 2016; Geddes et al., 2018).

The goal of the study by Cohen and colleagues (2016) was to examine possible age-related differences in neural recruitment during the presentation of high and low point values (i.e., reward anticipation phase) that preceded the to-be-remembered words (i.e., target stimulus phase). In line with previous behavioral studies detailed above, both younger and older adults showed greater recall for high-value words compared to low-value words, but the fMRI data revealed age differences in the underlying neural patterns. Activity in the reward network during the reward anticipation phase was modulated by reward value in younger adults, but not older adults, and in neither group was this activation associated with value-based memory selectivity. These neural findings are in contrast to other work with younger adults showing that engagement of the reward network during the reward anticipation phase leads to greater memory for the subsequent to-be-remembered stimulus (e.g., Adcock et al., 2006). Instead, Cohen and colleagues found that for younger adults, activity within the reward network during this later target stimulus phase correlated with value-directed memory selectivity, but not in older adults.

In the study by Geddes and colleagues (2018), older and younger adults completed the MIE task while undergoing fMRI. In contrast to the behavioral studies detailed above (Castel et al., 2002; Cohen et al., 2016; Mather and Schoeke, 2011; Spaniol et al., 2014), older adults did not show evidence of reward motivation effects on behavioral memory performance. Only younger adults demonstrated modulation of memory performance as a function of reward motivation (both gain and loss), and this effect was limited to recollection-based (rather than familiarity-based) memory. Similar to the findings reported by Cohen and colleagues (2016), activation in the reward network during the target stimulus phase was associated with motivation-related memory gains, but this effect was limited to younger adults. Both age groups activated the reward network as well as memory related regions during anticipation of gains and losses, but this activity was not correlated with memory performance in either group.

In sum, the two existing fMRI studies on aging and reward-modulated memory arrived at contradictory conclusions. Cohen and colleagues replicated the behavioral effect of reward-modulated memory, but activity in the reward network during the target stimulus phase, not the anticipation phase, predicted behavioral performance, and only in younger adults. Geddes and colleagues did not replicate these behavioral findings in older adults even though older adults showed intact anticipatory responses to reward cues. There were many differences between these two studies that could potentially account for the discrepant findings. For example, differences in the type of motivation (i.e., points vs. monetary rewards), in motivation valence (Geddes and colleagues included both gains and losses, whereas Cohen et al. only examined gains), retention interval (immediate vs. delayed), and how the reward network was defined for analyses (i.e., derived from neurosynth.org vs. functional localizer from the participants).

### The Current Study

In the current study, younger and older adults completed the MIE task during fMRI scanning, similar to Adcock et al. (2006) and Geddes et al. (2018). However, we analyzed neural activity from a different perspective than previous studies. In addition to probing activation within regions of the reward network, we also examined the functional connectivity between reward-related regions and the rest of the brain, during both the reward anticipation and the target presentation phases. We hypothesized that we would replicate our prior findings from a separate behavioral experiment of reward-based memory enhancement in the MIE task in both age groups (Spaniol et al., 2014). Further, based on our previous observations in the context of an incentive processing task with the same participants included in the current study (Spaniol et al., 2015), we hypothesized that in addition to the reward network, older adults would also show increased engagement of prefrontal regions in the service of reward-enhanced memory. Whether this pattern would occur during the reward anticipation phase indicative of preparatory, proactive cognitive control that assists in the maintenance of goal information, or during the target stimulus phase indicative of delayed, reactive control (Braver, 2012; Chiew and Braver, 2014; Fröber and Dreisbach, 2014) was an open question given mixed findings in the literature (Cohen et al., 2016; Geddes et al., 2018).

## Method

### Participants

Ethics approval for all procedures was obtained from Baycrest Hospital and Ryerson University. Younger adults were recruited from Ryerson University and the Toronto area via community websites (Craigslist.ca and Kijiji.ca) and older adults were recruited from both the Baycrest Hospital and Ryerson University older adult participant pools. Study eligibility was established prior to scheduling, and determined with a screening questionnaire to assess past and current medical conditions (e.g., psychiatric illness, depression, head injury, stroke), and medications that may affect cognition (e.g., sleep aids, prescription pain medication). Contraindications to the MRI procedure (e.g., metal implants) were assessed with an MRI safety questionnaire. Sixteen young adults (9 females) and 17 older adults (9 females) completed the study. Two older adult males were excluded from all analysis; one due to an incidental MRI finding, and one for failing to follow instructions. One additional older adult male was excluded from the connectivity analyses due to extreme outlier percent signal change values in the brain data, leaving a final sample of 14 older adults. Participants were compensated $80 for participation over the two sessions, in addition to task-based earnings.

Older adults were 67.8 years old on average (SD = 4.69, range = 60-76) and younger adults 25.44 (SD = 3.79, range = 20-33) years old. All older adults scored 27 or higher on the Mini Mental State Exam (Folstein et al., 1975), with the exception of one with a score of 26, but this individual was not excluded from the analyses since no other measures showed evidence of impairment, and excluding the participant did not change the pattern of results. The two groups did not differ on years of education (M_older_ = 16.5, SD = 1.95; M_younger_ = 16.69, SD = 2.85), nor on any subscales of the revised 60-item NEO Five-Factor Inventory (Costa and McCrae, 1989). Older adults scored higher than younger adults on the positive mood scale (M_older_ = 33.29, SD = 7.30; M_younger_ = 28.25, SD = 5.75) of the Positive and Negative Affect Schedule (PANAS; Watson etal.,1988), *t*(28) = 2.11, *p* = .04, η^2^ = .13, and on the Mill-Hill vocabulary scale, *t*(27)^1^ = 3.91, *p* = .001, η^2^ = .35 (M_older_ = 23. 46, SD = 3.77; M_younger_ = 18.38, SD = 3.22).

### Stimuli

The experimental stimuli consisted of 240 color photographs consisting of 120 indoor scenes and 120 outdoor (see Figure 1 for examples), taken from a picture database in CorelDraw. Stimuli were devoid of people or animals. The scenes were divided into four sets of 60 stimuli (half indoor, half outdoor). The sets contained approximately equal proportions of specific types of indoor scenes (e.g., kitchens, living rooms) and outdoor scenes (e.g., deserts, mountains). Sets of stimuli were counterbalanced across reward value and participants. An additional 12 stimuli were used for the practice runs.

**Figure 1.**
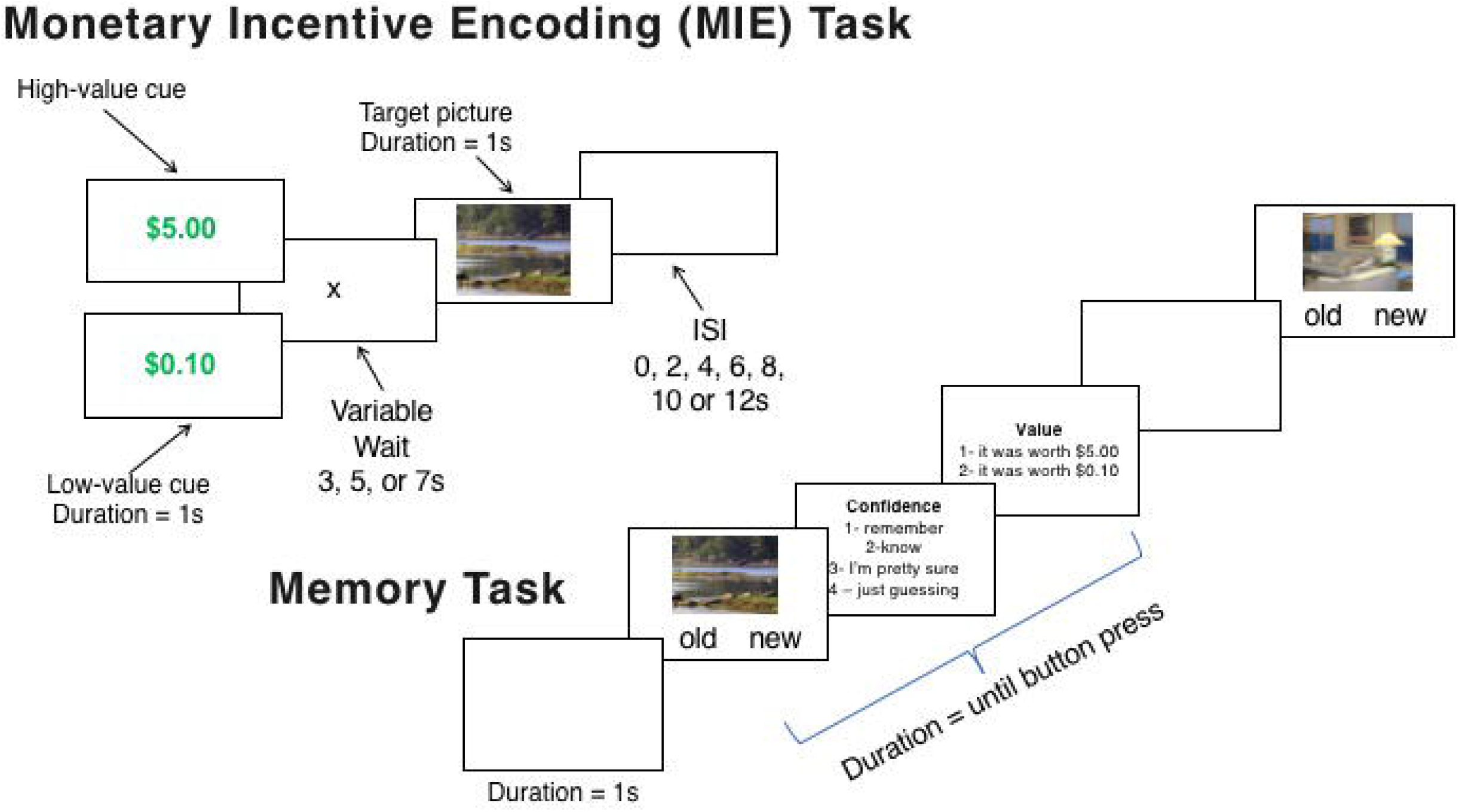
Trial procedure in the Monetary Incentive Encoding (MIE) Task and recognition memory task. The MIE task was completed in the fMRI scanner and duration of each element of the procedure is reported. The memory task depicts a trial resulting in an “old” response followed by a confidence rating and value judgment.

### Paradigm

The paradigm closely followed the procedures from prior work (Adcock et al., 2006; Spaniol et al., 2014). While in the MRI scanner, participants completed three runs of the Monetary Incentive Encoding (MIE) task (Adcock et al., 2006). During this task, participants were asked to intentionally encode 120 scene stimuli (40 trials per run). The target images remained onscreen for 2,000 ms and were preceded by a reward cue. The reward cue indicated how much money the participant could earn if the image was correctly recognized on the memory test the following day. Cues either indicated a high value ($5.00) or low value ($0.10) and remained onscreen for 1,000 ms. The cue was followed by a fixation that remained onscreen for a variable length of 3,000, 5,000 or 7,000 ms. Optseq was utilized to create a randomized schedule of between trial interstimulus interval lengths of 2,000, 4,000, 6,000, 8,000, 10,000, and 12,000 ms based on the study parameters (e.g., number of time points, repetition time). See Figure 1 for a depiction of the MIE task.

After completing the MIE task and while still in the MRI scanner, participants then completed a functional localizer task—the Monetary Incentive Delay (MID) task—well-established to localize the reward network of the brain (Dugré et al., 2018; Knutson et al., 2001b, 2000). We have described the details of this task in prior publications (Bowen et al., 2019; Spaniol et al., 2015)^2^, but briefly, a participant’s reaction time to a fast-paced target is incentivized with monetary gains and losses. Like the MIE task, each trial begins with a reward cue (i.e., reward anticipation phase) indicating how much money the participant can gain or avoid losing if the target stimulus—a star—is responded to in the allotted amount of time. The duration that the target remains on screen is calibrated to speed-up and slow-down so participants maintain a hit rate of approximately 66%.

The day after completing the MIE task, participants completed a recognition test with the 120 stimuli previously encoded in the MRI scanner and 120 new stimuli. For each image participants were asked to first indicate old or new, if new was selected the next image was presented. If old was selected, participants were asked to rate their confidence in their memory for the image on a scale of 1 to 4. Participants were told to respond ‘1’ if “you remember the moment that you encountered the picture”; respond ‘2’ if “you feel sure that the picture was presented, but you have no specific memory”; respond ‘3’ if “you are pretty sure, but not certain, that the picture was old’; respond ‘4’ if “you were just guessing”. Following the confidence rating, participants were asked a value judgment and to indicate whether they thought the image was worth a high ($5.00) or low reward ($0.10) value. See Figure 1 for a depiction of the recognition memory task.

During the recognition test, if participants correctly identified an image as “old”, they earned either the $5 or $0.10 the image was worth, however if they incorrectly identified a new image as old, they lost $2.55. This penalty for false alarms was included to reduce liberal responding on the memory test.

### Procedure

#### fMRI and Behavioral Procedures

Session 1 took place at Baycrest Hospital where participants were met by the researcher. After obtaining informed consent, participants changed into scrubs and spent the first 30 minutes reading instructions and practicing the MIE task. Practice took place in the MRI simulator to ensure that participants were comfortable with the MRI environment and the necessary responses and button presses. After setup in the MRI, the T1 anatomical scan was collected followed by the MIE task. This was an event-related design and participants completed three runs of forty trials, each run was 7 minutes and 36 seconds. The MIE task was followed by the MID task. After completion of the MRI session, participants changed out of scrubs, were paid for their participation that day and earnings on the MID task, and reminded of their appointment the following day.

Session 2 took place approximately 24 hours after encoding, at Ryerson University. Participants completed the questionnaires followed by the recognition task described above which took approximately 40 minutes. At the end of the session participants were paid the amount they had earned for participation on that day, in addition to the performance-based rewards they earned on the MIE memory test.

#### fMRI data acquisition

MRI scanning was conducted on a Siemens Trio 3.0T whole-body scanner using a 32-channel “matrix” head coil. Anatomical imaging protocol included three dimensional T1-weighted imaging (MPRAGE, FOV = 25.6 cm^2^, 1×1×1 mm voxels, TI/TE/TR = 1100/2.63/2000 ms, flip angle = 9 deg, averages = 2, 160 slices, scan time = 5:44) and fluid attenuated inversion recovery imaging (interleaved axial multislice FLAIR, FOV = 22.4 cm^2^, 0.9×0.9×5mm voxels, bandwidth = 315Hz/Px, TI/TE/TR = 2200/96/9000ms, averages = 1, concatenations = 3, 32 slices, 5mm thickness, scan time = 3:38). Functional (MIE task) scans were acquired using an interleaved multislice EPI sequence (oblique axial orientation intercallosal line, 228 volumes; FOV = 19.2 cm^2^, 64×64 acquisition matrix, 40 slices 3 x 3 x 3.5 mm in-plane resolution, bandwidth = 2604 Hz/Px, TE/TR = 27/2000 ms, flip angle = 70 deg). Stimulus presentation and image acquisition was synchronized with a trigger pulse sent by the scanner at the beginning of each experimental run. Using an LCD projector (NEC Model MTI065) with a 2.75-5 zoom lens (Navitar, Inc.), visual stimuli were projected on a screen at the back of the magnet bore and viewed by the participant through a mirror attached to the head coil. Responses to the stimuli were made via a Fiber-Optic Response Pad System (Current Designs Inc.; 4 buttons available per hand). fMRI-compatible prescription glasses were available to correct for visual acuity (SafeVision LLC., −6 to +6 diopters available in 0.5 increments). To reduce movement, foam sponges were used to restrain the participant’s head and physiological data (heart rate, respiration, pulse) were also collected. To allow magnetic stabilization, the first 4TRs of the functional runs were discarded.

### fMRI Data Analysis

#### Analysis logic

Overall, we were interested in both activation in a region of the reward network, and functional connectivity patterns between that region and the rest of the brain. We examined these neural patterns in both younger and older adults during subsequent memory for high and low reward items. To this end, we first established an ROI in the reward network using an activation analysis from the functional localizer MID task. From this activation analysis, the caudate—a well-established region of the reward network—was chosen as the ROI. Activation patterns with this caudate seed were examined during the anticipation and target stimulus phase of the MIE task. A whole-brain generalized psychophysiological interaction (gPPI) functional connectivity analysis using the caudate as a seed was conducted at the anticipation phase of the MIE task. These results are presented below.

#### Pre-processing of fMRI data

Using SPM12 software (Wellcome Department of Cognitive Neurology, London, United Kingdom), pre-processing steps of functional images included reorienting realignment, as well as coregistration of image volumes to a standard MNI-space template, resampling to 2mm^3^ voxels (EPI.nii) and spatial smoothing with a 4-mm Gaussian kernel. To detect motion and global mean intensity outliers we used Artifact Detection Tools (ART; available at www.nitrc.org/projects/artifactdetect). The parameters for outlier detection were the following: 1) more than 3 standard deviations above the global mean intensity, 2) less than ±5 mm translation motion and 3) ±1 degree rotation. A run was considered problematic if more than 10TRs were detected as outliers. No run or trials were removed for any participants based on these criteria. These pre-processing steps and artifact detection parameters were used for both the MIE and MID task.

### MID Task

#### GLM

The functional localizer MID task was used to find an ROI within the reward network. After the data were pre-processed, at the first (subject) level of analysis, an event-related design matrix was created. It was modelled around the onset of the reward cue, contained 4 columns of interest (High Gain, Low Gain, High Loss, Low Loss) and 3 columns to regress out linear drift for the three concatenated runs. Contrasts were then created comparing each condition of interest to baseline (e.g., High Gain Cue > Baseline). A full factorial ANOVA was used at the second (group) level to examine activation during the reward anticipation phase of the experiment. Reward (high, low) and valence (gain, loss) were entered as within-subjects variables and age (young, old) was entered as a between-subjects variable.

#### ROI

To identify an ROI to be used in subsequent analyses, we used the results from the GLM of the MID task. In line with prior work (Adcock et al., 2006) activation during the High Gain > Low Gain cue (across young and older adults) was used to isolate the reward network. This analysis revealed widespread activity in the mesolimbic reward network (see Figure 2A) with the most significant cluster in the left caudate (MNI: X = −8, Y = 6, Z = 2), a region that has previously been implicated as important for reward processing (Knutson et al., 2001a; Wilson et al., 2018; Zink et al., 2004). Using the SPM toolbox MarsBaR (Brett et al., 2002) a 10 mm sphere around the left caudate was used to create an ROI (see Figure 2B) to be used in the subsequent analyses with the MIE task, detailed next.

**Figure 2.**
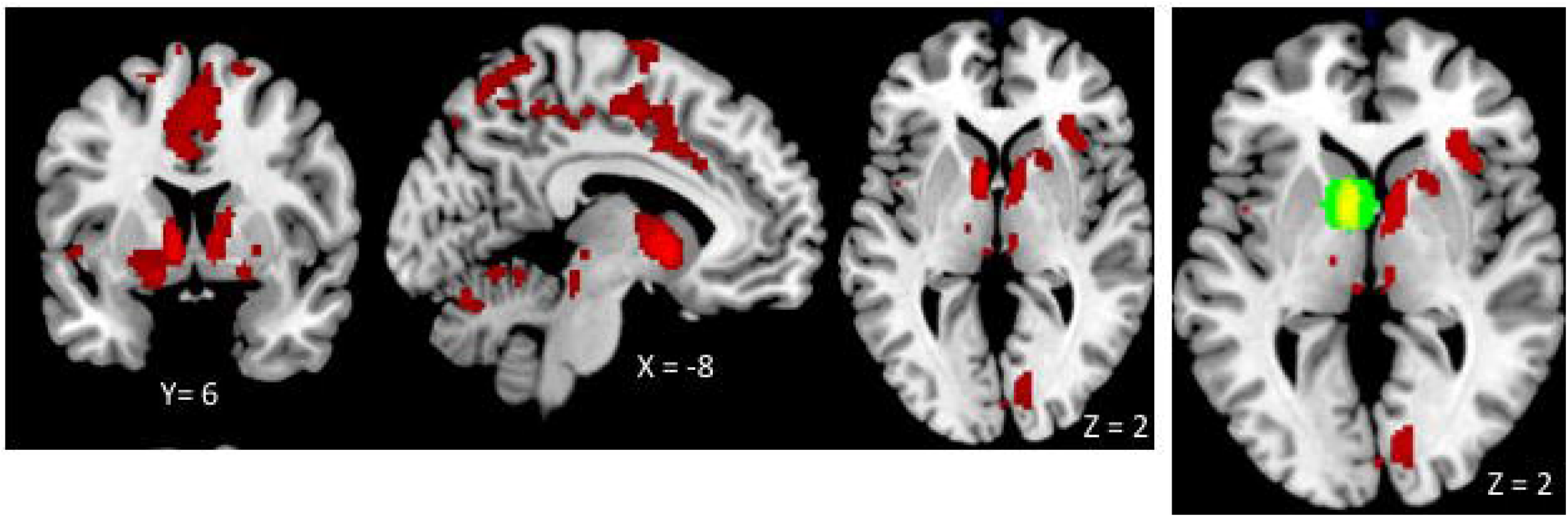
A) Depicts the activation from the Monetary Incentive Delay (MID) task during the reward anticipation (cue) phase with the contrast High Reward > Low Reward across younger and older adults. B) The green circle indicates the 10mm sphere created around the left caudate (MNI: X = −8, Y = 6, Z = 2) to use as an ROI in all other analyses.

### MIE Task

#### Activation Analysis

In line with prior work examining activation associated with reward-modulated memory in younger adults (Adcock et al., 2006), the current fMRI analysis of the MIE task contains only high confidence trials, combining “remember” and “know” responses. At the first level of analysis, two event-related design matrices were created. One was modelled around the onset of the reward cue, and the second around the onset of the target stimulus. For both models, an 8-column regression matrix included 4 conditions of interest (subsequent hit and subsequent miss for both high and low reward), 1 nuisance regressor column of low confidence trials, and 3 columns to regress out linear drift for the three concatenated runs. Contrasts were then created comparing each condition of interest to baseline (e.g., Subsequent High Reward Hits > Baseline). Consistent with prior work (Adcock et al., 2006; Cohen et al., 2016; Geddes et al., 2018), activation within the reward-related region was examined as a function of age, memory status, and reward. Specifically, activation within the caudate ROI created from the MID task (described above) was extracted from each subject’s first level models using the REX toolbox (downloaded from http://web.mit.edu/swg/software.htm) and entered into a mixed ANOVA. See supplementary material for the whole-brain activation analyses.

#### Connectivity Analysis

The activation analysis described above was used to determine whether age influenced the extent to which a reward-related region preferentially supported memory for high relative to low reward items. When age invariance was identified, follow-up connectivity analyses were conducted to determine whether this reward-related region supported memory for high versus low reward items in the same way for young and older adults. These connectivity analyses thus probed for possible age-variant networks supporting high compared low reward memory through their connections with an *age-invariant* reward region. The generalized psychophysiological interactions (gPPI) toolbox (http://brainmap.wisc.edu/PPI; McLaren et al., 2012) was used to compare functional connectivity with a single seed region across task conditions. Specifically, we examined functional connectivity between the caudate ROI and the rest of the brain during the presentation of the reward cues of the MIE task. First, a 6 mm volume of interest (VOI) for each subject was created around the caudate ROI peak (MNI: X = −8, Y = 6, Z = 2). Within each subject, the gPPI toolbox estimated the functional connectivity across the entire brain with this VOI in the 5 memory conditions (high reward hits, high reward miss, low reward hit, low reward miss, low confidence hits) to calculate the 4 contrasts of interest compared to baseline. At the group level, the individual subject gPPIs were entered into full-factorial ANOVAs with reward (high, low), memory (hit, miss) as within-subjects factors and age as a between-subjects factor. Voxel threshold for these analyses was set at *p* < 0.005 (uncorrected), Monte Carlo simulations (https://www2.bc.edu/sd-slotnick/scripts.htm) run with the normalized voxel size of 2 x 2 x 2, determined that a 17-voxel extent (k) corrected results to p < .05. Activation within the caudate ROI created from the MID task (described above) was extracted using the REX toolbox (downloaded from http://web.mit.edu/swg/software.htm). MNI coordinates from SPM12 were converted to Talairach Coordinates using GingerAle (http://www.brainmap.org/ale). Anatomical labels in the cluster report tables were assigned using the Talairach Daemon (Lancaster et al., 1997) and visually checked using an anatomy atlas (Talairach and Tournoux, 1988).

## Results

### Behavioral Results

Reaction-time (RT) results from the MID functional localizer task have been reported in prior publications (Bowen et al., 2019; Spaniol et al., 2015). Behavioral data from the MIE task were analyzed using SPSS (Statistical Package for Social Sciences 24.0. SPSS Inc., Chicago, USA). In line with prior work (e.g., Adcock et al., 2006) the results reported below are restricted to high confidence trials only (i.e., collapsed across “remember” and “know”) and exclude low confidence trials (i.e., “pretty sure” or “guessing” responses).

#### Reward earnings

On average, younger adults earned higher memory-based rewards (*M* = $101.17, *SE* = $15.10) than older adults (*M* = $64.91, *SE* = $7.30), *t*(28) = 2.07, *p* = .05, η^2^ = .13.

#### RT for recognition hits

Median reaction times for correctly recognized items (i.e., recognition hits) were submitted to a 2 (Reward Magnitude: high [$5], low [$.10]) x 2 (Age: younger, older) mixed ANOVA. There was no main effect of either variable, nor a significant interaction, *F*(1, 28) ≤ 1.96, *p* ≥ .17, η_p_^2^ ≤ .07. The average median reaction time across groups and conditions was 2,525 ms (*SE* = 159 ms).

#### Memory accuracy

Hit rates were submitted to a 2 (Reward: high, low) x 2 (Age: younger, older) mixed ANOVA. There was a main effect of reward, *F*(1, 28) = 9.98, *p* = .004, η_p_^2^ = .26, such that recognition was better for high-reward items (*M* = .56, *SE* = .04) compared to low-reward items (*M* = .47, *SE* = .04), but there was no main effect of age, *F*(1, 28) = .05, *p* = .82, η_p_^2^ = .002, nor a significant interaction, *F*(1, 28) = .49, *p* = .49, □_p_^2^ = .02. Since new items were never paired with a reward value there was only a single false alarm rate which did not differ significantly for younger adults (*M* = .13, *SE* = .03) and older adults (*M* = .20, *SE* = .03), *t*(28) = 1.69, *p* = .10, η_p_^2^ = .09.

#### Memory confidence

To examine whether older and younger adults differed in their use of “high” versus “low” confidence levels for recognition hits, the proportion of high and low confidence levels were submitted to a 2 (Confidence Level: high, low) x 2 (Reward: high, low) x 2 (Age: younger, older) mixed ANOVA. There was a significant Confidence Level x Age interaction, *F*(1, 28) = 9.69, *p* = .004, η_p_^2^ =.26, and a significant Confidence Level x Reward interaction, *F*(1, 28) = 7.74, *p* = .01, η_p_^2^=.22. No other interactions nor main effects were significant, *F*(1, 28) ≤ 2.20, *p* ≥ .15, η_p_^2^ ≤ .07. Breaking down the first interaction with follow-up independent samples *t*-tests, younger adults used high confidence more often for hits (*M* = .28, *SE* = .03) than older adults (*M* = .18, *SE* = .02), *t*(28) = 3.07, *p* = .005, □^2^ = .25. Older adults used low confidence more often for hits (*M* = .32, *SE* = .02) than younger adults (*M* = .22, *SE* = .02), *t*(28) = 3.16, *p* = .005, □^2^ = .26. Follow-up paired sample *t*-tests for the second interaction revealed no significant simple main effects. Participants did not differ in their use of high (*M* = .22, *SE* = .02) and low (*M* = .28, *SE* = .02) confidence for high-reward items, *t*(29) = 1.39, *p* = .17, □^2^ = .06, nor in their use of high (*M* = .24, *SE* = .02) and low confidence (*M* = .26, *SE* = .02) for low-reward items, *t*(29) = .46, *p* = .65, □^2^ = .01.

#### Memory for reward value

Accuracy of reward value judgements was calculated for correctly identified target items. Hit rates were submitted to a 2 (Reward: high, low) x 2 (Age: younger, older) mixed ANOVA. There was a significant main effect of reward, *F*(1, 28) = 9.98, *p* = .004, η_p_^2^ ≤ .26, such that value accuracy was higher for high-reward (*M* = .56, *SE* = .04) compared to low-reward (*M* = .47, *SE* = .04) items. No other effects were significant, *F*(1, 28) ≤ .49, *p* ≥ .49, η_p_^2^ ≤ .02.

### fMRI Results

To align with the behavioral analyses, the fMRI analyses reported below include only high-confidence trials. We first report the results of activation within the caudate ROI followed by the gPPI functional connectivity analysis. Whole-brain activation analyses are presented in the supplementary material.

#### Activity in the ROI during MIE task

Activity within the 10mm sphere created around the caudate (MNI: X = −8, Y = 6, Z = 2) was extracted at anticipation phase and the target stimulus phase of the MIE task. Activity (reported as percent signal change) from these phases was then entered into separate repeated-measures ANOVAs with reward (high, low) and subsequent memory (hit [recognized], miss [forgotten]) as within-subjects variables and age (young, old) as a between-subjects variable.

##### Anticipation phase

There was a main effect of reward, *F*(1, 29) = 6.41, *p* = .02, η_p_^2^ = .18, qualified by a Reward x Memory interaction, *F*(1, 29) = 8.53, *p* = .007, η_p_^2^ = .23. Follow-up paired samples t-tests indicated a subsequent memory effect (hits > miss) for high-reward, but not low-reward items. Activity was greater for subsequently recognized (*M* = .40, *SE* = .11) compared to subsequently forgotten high-reward items (*M* = .09, *SE* = .08), *t*(30) = 2.38, *p* = .02, η^2^ = .16 There was no difference in activity within the caudate for subsequently recognized (*M* = −.09, *SE* = .13), compared to subsequently forgotten (*M* = .09, *SE* = .08) low-reward items, *t*(30) = 1.55, *p* = .13, η^2^ < .07. No other main effects nor interaction were significant, including no significant effects of age, *F*(1, 29) ≤ 2.60, *p* ≥ .12, η_p_^2^ ≤ .08.

##### Target phase

There were no significant main effects of reward, subsequent memory, or age, and no significant interactions, *F*(1, 29) ≤ 3.11, *p* ≥ .09, η_p_^2^ ≤ .09.

##### gPPI

The ROI results presented above indicate an age-invariant role of the caudate in supporting memory for high-reward items during reward anticipation. However, it is unclear from this activation analysis whether the caudate supports high-reward memory *in the same way* in young and older adults. A functional connectivity analysis was conducted to identify regions preferentially associated with the caudate prior to successful encoding of high compared to low-reward items. Further, we examined whether the networks associated with caudate activation would differ as a function of age, despite the age invariance of the caudate activation itself. In the current paradigm, during the target stimulus phase the caudate did not exhibit greater subsequent memory effects for high versus low reward items, as would be expected if it was contributing to reward-related memory enhancements. Because there is no evidence that participants, regardless of age, relied on this region to enhance high reward memory performance during the target phase, it was not necessary to examine how additional regions might be recruited during this phase to support reward-related memory enhancement. As a result, we only performed functional connectivity analyses for the anticipation phase.

Critically, the gPPI analysis revealed a significant Reward x Memory x Age interaction in connectivity between the caudate and several regions, including bilateral inferior frontal gryus, pre-and postcentral gyrus, superior parietal lobule, medial temporal gyrus, cingulate gyrus, precuneus, cuneus, inferior occipital gyrus, and insula. See Table 1 for all regions that showed effects of reward, memory, and age on caudate connectivity and supplementary materials Figure 3 for a visual depiction of the neural pattern. To better understand the direction of the three-way interaction, we extracted beta estimates from a theoretically-motivated subset of regions. We were interested in areas of the frontal lobe to test our hypothesis that older adults would show increased engagement of prefrontal regions in the service of reward-enhanced memory and contrast this with the pattern in occipital regions as several studies of neurocognitive aging have shown an age-related reduction in posterior neural activity and increased frontal activity (Davis et al., 2008; Grady et al., 1994; Li et al., 2015), often referred to as the posterior-anterior shift in aging (PASA).

**Figure 3.**
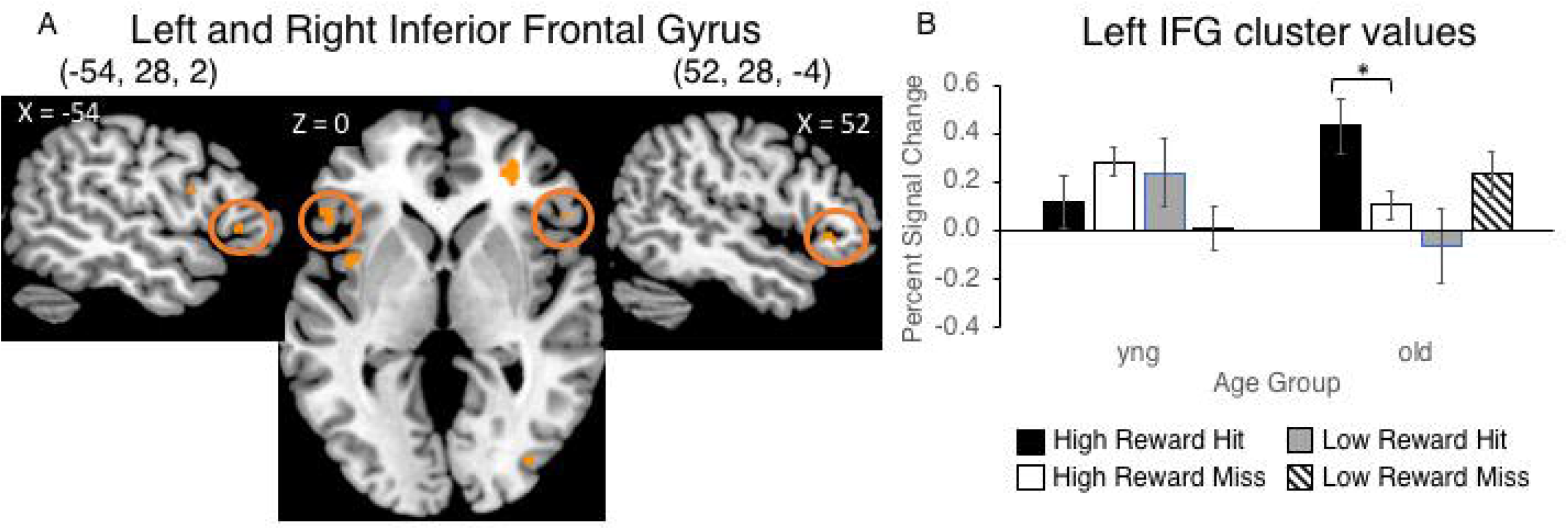
A) Left and right inferior frontal gyrus showing a three-way interaction in connectivity with the caudate ROI. B) Graph of the percent signal change depicting the pattern of connectivity of the three-way interaction. Values in the graph are from the left IFG cluster, but the pattern is the same for right IFG. Yng = younger adults; old = older adults.

**Table 1.**
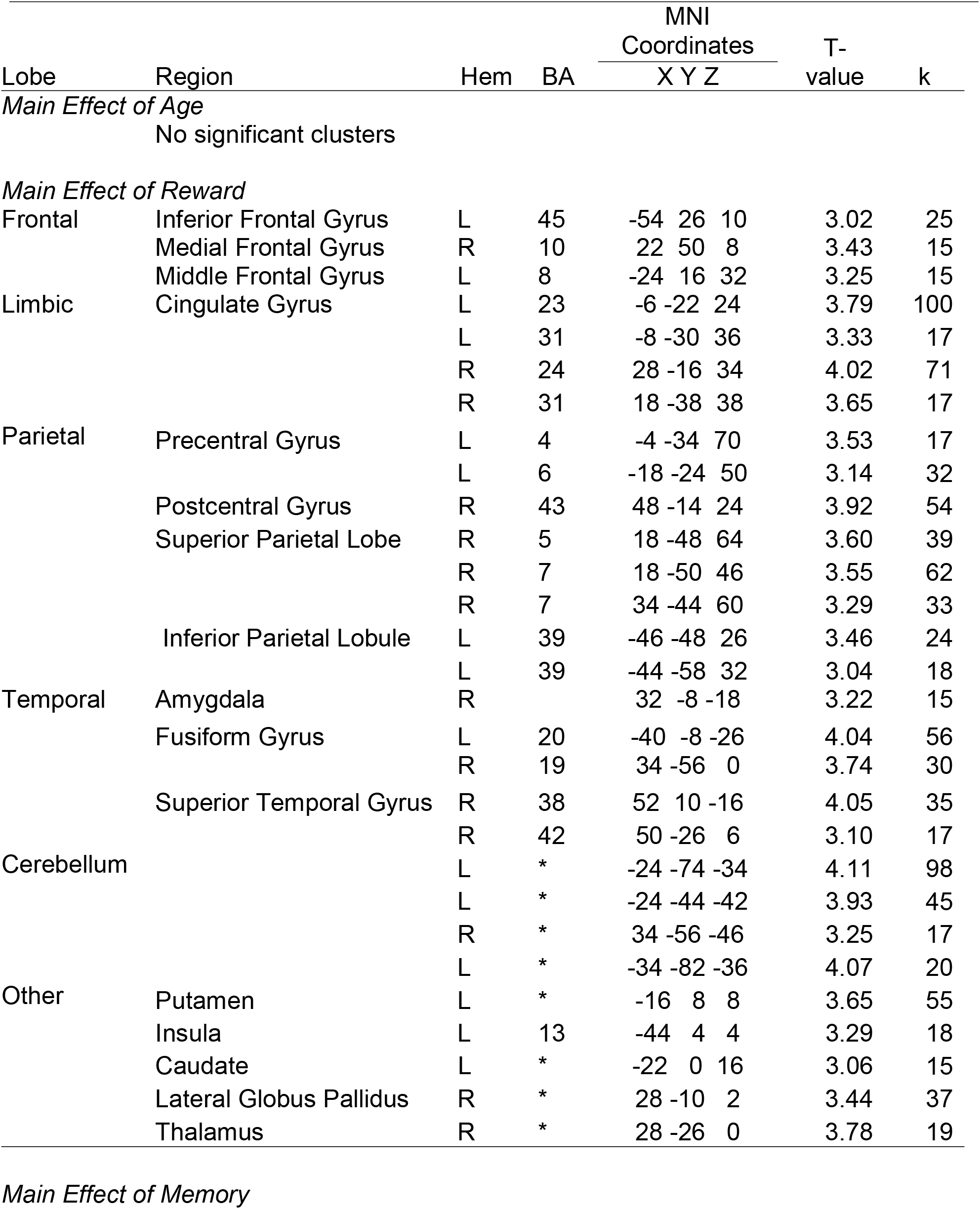
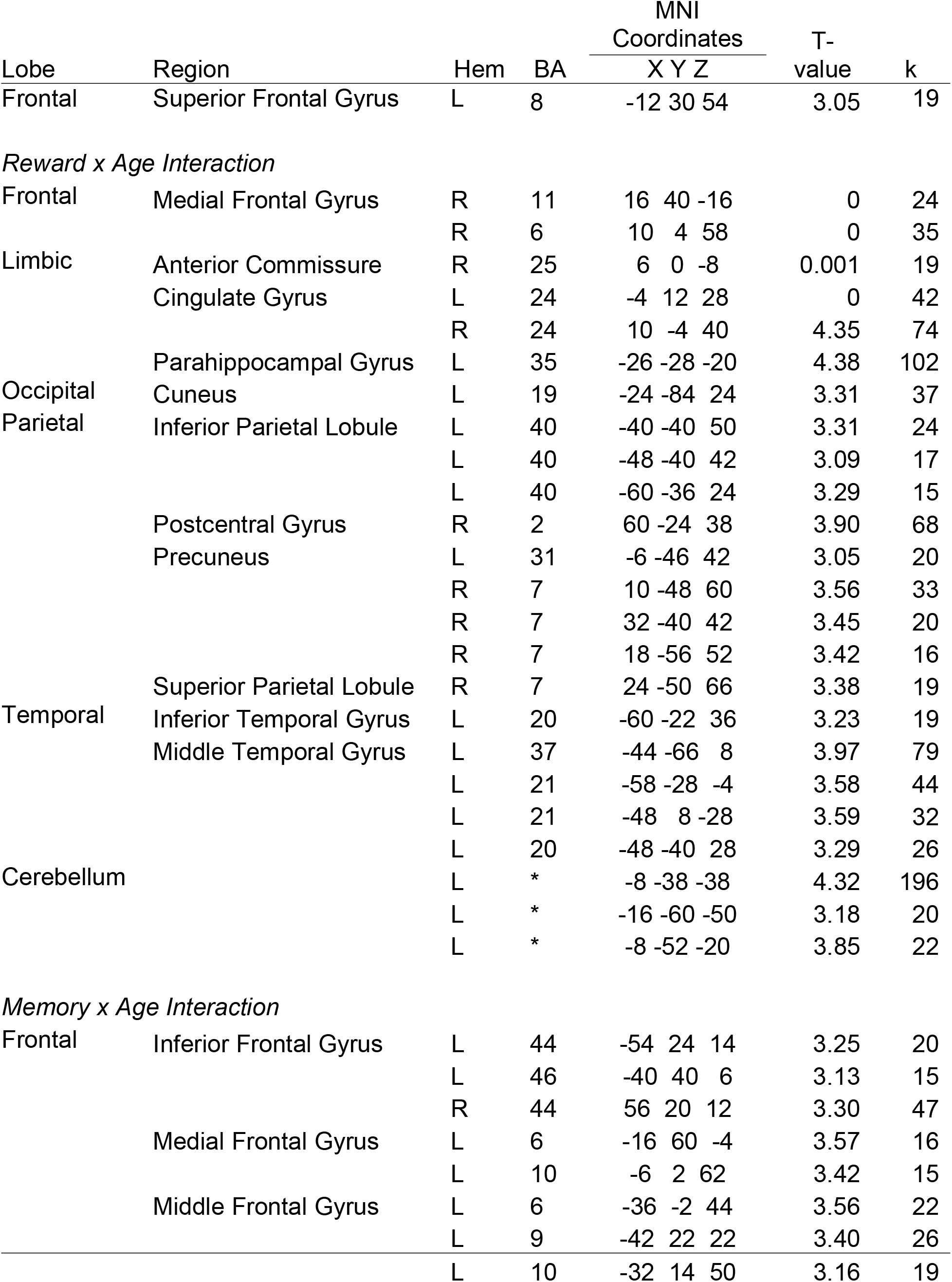
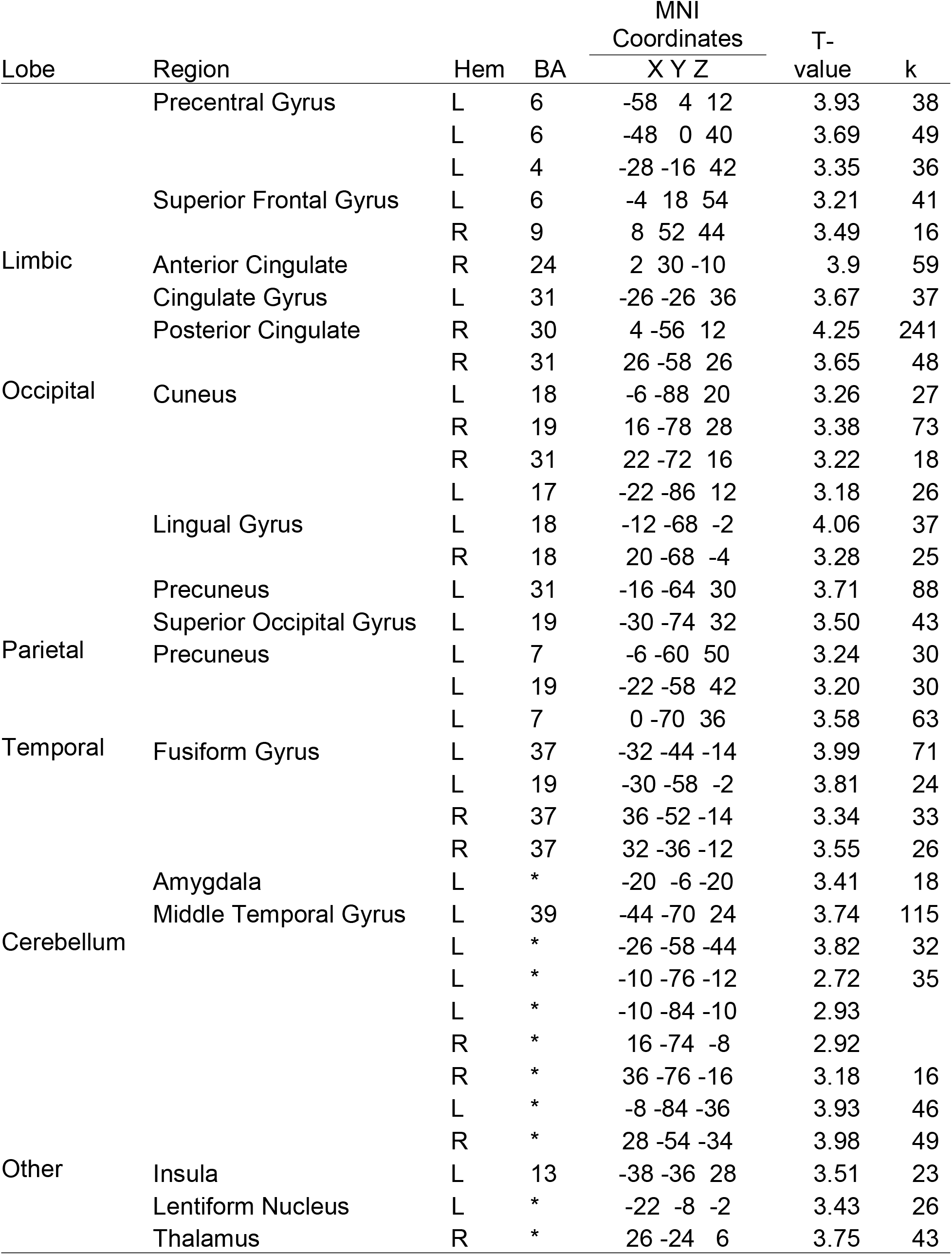
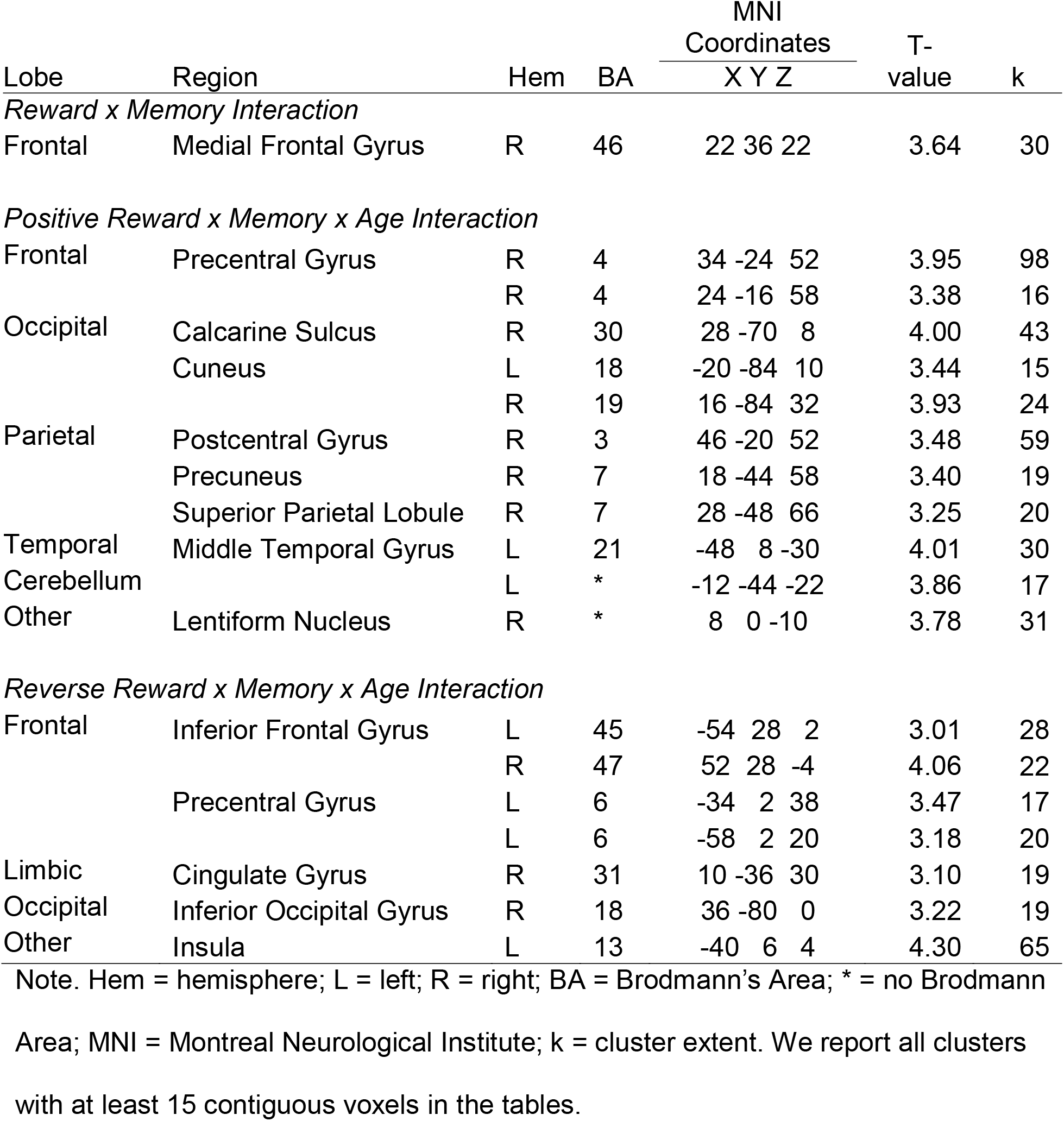
*Regions exhibiting differential neural connectivity with the caudate as a function of reward, aging and subsequent memory*

The three-way interaction with the left IFG was driven by significant Reward x Memory interaction for both younger adults, *F*(1, 15) = 5.72, *p* = .03, η_p_^2^ = .28, and older adults, *F*(1, 13) = 4.71, *p* = .05, η_p_^2^ = .27. In younger adults, connectivity between these regions did not differ for subsequently recognized versus forgotten high-reward items, *t*(15) = 1.54, *p* = .14, η^2^ = .14, but connectivity was marginally greater for subsequently recognized compared to forgotten low-reward items, *t*(15) = 1.85, *p* = .08, η^2^ = .19. For older adults, connectivity between the caudate and left IFG was significantly greater for subsequently recognized versus forgotten high-reward items, *t*(13) = 2.47, *p* = .03, η^2^ = .32, but not for subsequently recognized compared to forgotten low-reward items, *t*(13) = 1.31, *p* = .21,η^2^ = .12. The right IFG did not show a Reward x Memory interaction for younger adults, nor any other effects, *F*(1, 15) ≤ 3.11, *p* ≥ .10, η_p_^2^ ≤ .17. For older adults, the Reward x Memory interaction was significant, *F*(1, 13) = 13.0, *p* = .003, η_p_^2^ = .50. This interaction was driven by a marginally significant subsequent memory effect in connectivity between these regions for high-reward items, *t*(13) = 2.04, *p* = .06, η^2^ = .24, but significantly less connectivity between these regions during subsequently recognized compared to forgotten low-reward items, *t*(13) = 2.17, *p* = .05, η^2^ = .27. See Figure 3A for depiction of the IFG brain regions and Figure 3B for graphical representation of this pattern.

The three-way interaction with posterior sensory regions including the precuneus and the cuneus was driven by a pattern different from the one seen in the frontal regions. The interaction in the precuneus and the cuneus was driven by a main effect in younger adults, *F*(1, 15) ≥ 4.02, *p* ≤ .06, η_p_^2^ ≥ .21, such that there was greater connectivity between these regions and the caudate during high-reward cues compare to low-reward cues. For older adults, the Reward x Memory interaction was significant, *F*(1, 13) ≥ 5.57, *p* ≤ .04, η_p_^2^ ≥ .30. This interaction was driven by decreased connectivity between these regions and the caudate during subsequently recognized compared to forgotten high-reward items, *t*(13) ≥ 2.35, *p* ≤ .04, η_p_^2^ = .30. There was no subsequent memory effect for low-reward items, *t*(13) ≤ 1.66, *p* ≥ .12, η_p_^2^ = .17. See Figure 4A for a depiction of the brain regions and Figure 4B graphical representation of this pattern for the left caudate seed.

**Figure 4.**
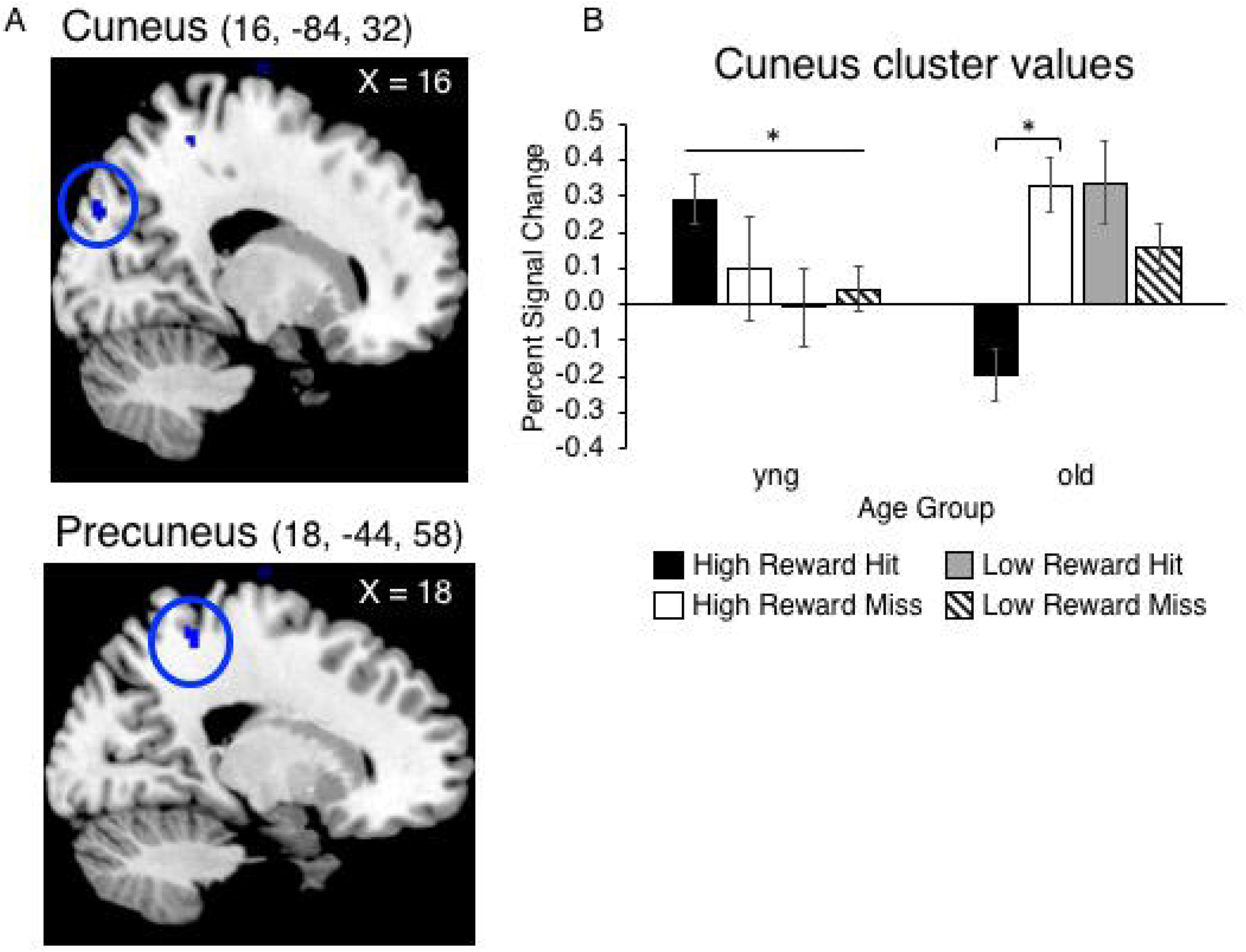
A) Cuneus (top) and precuneus (bottom) clusters that showed a three-way interaction in connectivity with the caudate ROI. B) Graph of the percent signal change depicting the pattern of connectivity of the three-way interaction. Values in the graph are from the right cuneus cluster, but the pattern is the same for all three regions. There was a main effect of reward in younger adults, and a Reward x Memory interaction for older adults. Yng = younger adults; old = older adults.

## Discussion

In the current study we sought to examine neural activation and functional connectivity patterns associated with reward-enhanced memory in healthy younger and older adults. In line with our hypothesis, high-reward items were better remembered than low-reward items by both age groups. Furthermore, both age groups showed greater activity in the left caudate, a reward-network region, during high-reward cues preceding subsequently remembered versus subsequently forgotten items. In line with our second hypothesis, functional connectivity between the caudate and lateral frontal gyri was associated with reward-enhanced memory in older adults. These results provide further evidence of a preserved influence of reward on memory formation in healthy aging, but they also point toward age differences in the nature of the link between reward and memory. In particular, increased fronto-striatal connectivity during reward cues that precede the presentation of target stimuli during encoding suggests that older adults engage proactive cognitive control processes to bolster memory for reward-related information. The finding also aligns with the observation of a posterior-anterior shift in aging (PASA; Davis, Dennis, Daselaar, Fleck, & Cabeza, 2008), and provides the first evidence of this pattern in the context of motivation-cognition interactions.

### Reward-Motivated Memory in Younger and Older Adults

Compared with younger adults, older adults earned lower performance-based rewards in the MIE task. However, like younger adults, older adults demonstrated better memory for items associated with high-reward compared to items associated with low-reward. This replicates findings from our two previous experiments with a different group of younger and older adults that used this same paradigm (Spaniol et al., 2014) as well as other findings of preserved value-based memory selectivity in aging (Castel et al., 2002, 2011; Cohen et al., 2016; for a review see Castel, 2007). Evidence that healthy older adults are sensitive to motivational incentives fits with theories of aging postulating that reward processes do not show the same age-related decline as cognitive domains, but remain intact or even improve in healthy aging (Mather, 2016).

### Activity in Caudate Associated with Reward Motivated Memory

The behavioral results for younger and older adults were similar, but they do not explain *how* older adults are performing this task and whether they are using the same neural mechanisms as younger adults to support this behavior. Utilizing a functional task to localize the reward network, we found extensive activation in the dopaminergic midbrain and reward network, choosing the left caudate, part of the dorsal striatum, to create an ROI. To align with prior studies (Adcock et al., 2006; Cohen et al., 2016; Geddes et al., 2018) we first investigated the level of activation within the ROI during the anticipation and target stimulus phase of the MIE task. Activation within the ROI during the anticipation phase was greater for subsequently remembered compared to forgotten high-reward items, across both older and younger adults. This region was not differentially activated by memory status (hits vs. miss) for low reward items nor was activity in this region sensitive to the task conditions during the target stimulus phase, in either younger or older adults. Our results support Adcock et al. (2006) who found that activation in the ventral striatum during reward anticipation was associated with reward-motivated memory in younger adults. Only two prior studies have examined age differences in neural activation during reward motivated memory. Cohen and colleagues (Cohen et al., 2016) investigated activity in a much larger ROI of the reward network and found that activity was modulated by reward cue, but this did not predict value-directed memory selectivity. Instead, activity in the ROI during the target stimulus phase correlated with value-direct memory selectivity, but only in younger not older adults. Geddes and colleagues (2018) did not report effects of reward on memory in older adults, although they did find evidence that the reward network was engaged to the same extent by older and younger adults during incentive processing of the MID task, which replicates our prior neuroimaging findings (Spaniol et al., 2015).

### Frontal-Striatal Functional Connectivity Supports Reward Motivated Memory in Older Adults

The novel analysis in the current study was investigating age-related differences in functional connectivity associated with reward-motivated memory. Studies with older adults (Cohen et al., 2016; Geddes et al., 2018) have only examined reward and age effects on activation within *a priori reward* network ROIs, leaving how the reward network might connect to other areas of the brain to support reward-motivated memory an open question. One study with younger adults (Adcock et al., 2006) reported that greater connectivity between the ventral striatum and hippocampus during reward anticipation predicted superior memory performance.

This connectivity analysis focused on two *a priori* regions of interest, and did not encompass the rest of the brain. Based on our prior findings that older adults recruit a network of regions during incentive processing, including bilateral PFC (Spaniol et al., 2015), an area often associated with control processes (Braver, 2012), and in light of the evidence for a posterior-to-anterior shift in aging (PASA; Davis et al., 2008), we hypothesized that older adults would show greater connectivity between the reward network and frontal regions during reward-motivated encoding. Our connectivity analysis indeed supported this hypothesis with greater connectivity between the caudate and bilateral inferior frontal gyrus that was associated with high-reward items that were subsequently remembered versus forgotten. Connectivity between these regions did not differ as a function of task condition for younger adults. Regions in posterior sensory cortices also showed connectivity with the caudate, but the pattern underlying the interaction was very different. Greater connectivity between the caudate and precuneus and cuneus, was associated with high-reward items that were later forgotten versus remembered, akin to a negative subsequent memory effect (Kim, 2011) for older adults. For younger adults, in contrast, connectivity between the caudate and posterior sensory regions supported the processing of high-reward cues compared to low-reward cues, regardless of subsequent memory success. Again, these findings are in line with PASA (Davis et al., 2008), as older adults relied less on posterior regions and more on frontal regions. This may suggest that compared to younger adults, older adults’ engagement proactive cognitive control mechanisms supports reward-motivated memory.

Interestingly, a whole-brain activation analysis conducted by Geddes and colleagues during reward anticipation revealed activation in visual association cortex during the anticipatory cue period for motivational compared to neutral trials, which they propose reflects heightened visual attention in both age groups during the presentation of the gain and loss cues. Other work with younger adults (Murty, Tompary, Adcock, & Davachi, 2017) has indicated that increased post-encoding connectivity between visual cortex, ventral striatum and hippocampus predicted associative memory for high, but not low-reward items. The current study could not address post-encoding connectivity, but it appears that connectivity between the reward network and visual cortex during reward anticipation may actually hurt memory for high-reward items, at least for older adults.

### Comparing Findings from Three Studies Examining Neural Recruitment in Older Adults

The findings between the two published papers examining the neural correlates of reward-motivated memory in older adults do not align, and are also different from our findings in a number of ways. Both Cohen et al. (2016) and the current study replicated prior work indicating that older adults are sensitive to rewards and have better memory for items associated with a high reward compared to a low reward. In Cohen et al., these rewards were points and, in our study, monetary incentives. In Geddes et al. there were monetary gains, losses and neutral zero-dollar trials. It is possible that the presence of gain and loss trials had a different psychological effect on older adults when in the context of a declarative memory task. Other studies have also used the same three trial types albeit of smaller value (Mather and Schoeke, 2011) and found that memory for older adults was improved for gain over loss and neutral conditions on a test of incidental, not intentional memory encoding similar to the MID task.

The temporal characteristics of reward-network engagement and its link to memory performance also varied across the three studies. In the current study, caudate activation during the anticipation phase, but not during the target presentation, was linked to subsequent memory performance in both younger and older adults. In contrast, Cohen et al. (2016) observed activation in reward regions that was predictive of value-directed memory selectivity in older adults during the target phase only. The temporal differences between our findings and those reported by Cohen et al. (2016) may result from differences in task demands between the two paradigms. First, our study featured a longer encoding session, as well as a 24-hour delayed recognition test outside the scanner, whereas participants in Cohen et al.’s (2016) study recalled the words directly after encoding. These task differences may have encouraged different encoding strategies in younger and older adults. Geddes et al. (2018) did not find activation at either stage that supported reward-motivated memory in older adults, but also did not find a behavioral reward effect on memory in older adults. The preponderance of existing evidence from studies of reward-motivated memory in younger and older adults (Castel, 2007; Cohen et al., 2016; Mather and Schoeke, 2011; Spaniol et al., 2014) suggests that older adults’ memory is responsive to reward, potentially via the engagement of proactive control during memory encoding processes. The nonsignificant reward effect on older adults’ memory reported by Geddes et al.’s (2018) forms an exception, but given that reward effects on memory are typically small (e.g., η^2^ = .13-.28, Spaniol et al., 2014;), occasional null findings are not unexpected.

### Limitations and Future Directions

One of the limitations of the current study is its small sample size. However, it should be noted that the current findings replicate prior behavioral findings of better memory for high-reward, compared to low-reward stimuli (Adcock et al., 2006; Castel et al., 2002; Cohen et al., 2016, 2014, Shigemune et al., 2014, 2010; Spaniol et al., 2014). A second potential limitation is that the current study included only gain cues, not punishments or monetary losses. Whether valence plays a role in the behavioral, as well as neural recruitment is an important theoretical question. Prior work has shown that monetary gains and losses do not have the same effects on incidental episodic memory in younger adults (Bowen & Spaniol, 2017) or older adults (Mather and Schoeke, 2011), and that monetary gains may lead to better performance than punishments for younger adults in a spatial memory task (Murty, LaBar, Hamilton, & Adcock, 2011). Additionally, healthy aging is often associated with a motivational shift that can manifest as a positivity bias in attention and memory (Carstensen, 1995; Mather and Carstensen, 2005) and differences in neural processing of gain compared to loss anticipation (Samanez-Larkin et al., 2007) and gain and loss feedback (Bowen et al., *in press*). Geddes and colleagues did include both gain and loss trials in their paradigm, but did not observe effects on memory in older adults. Although speculative, a possible reason for this may be that the presence of loss cues undermines task effort in older adults. An empirical test of this question may help clarify some of the discrepant findings across studies.

### Conclusion

We examined reward-motivated memory in younger and older adults and the neural correlates of this effect. The current study used a paradigm resembling those used in two prior studies, but relied on a different analysis approach, by examining whole-brain functional connectivity with a region in the reward network. We found that although there were no age differences in seed-region activation levels during the reward anticipation phase, older and younger adults showed different functional connectivity patterns. For older adults, greater connectivity between the caudate and bilateral inferior frontal gyrus supported successful subsequent memory for high-reward items. Connectivity between the caudate and posterior regions was greater during high compared to low-reward anticipation for younger adults across remembered versus forgotten items, but for older adults connectivity decreased between these regions in support of successful encoding of items that followed a high-reward. These findings add to the growing literature on motivation-cognition interactions in healthy aging, and provide novel evidence of an age-related shift in the neural underpinnings of reward-motivated encoding.

## Footnotes

1 Due to time constraints during the experiment, one older adult did not complete the Mill-Hill vocabulary scale.

2 Participants in these two prior studies where we reported detailed results of the MID task are the same participants included in the current study where we report results of the MIE task.

