## Supplementary Material for "Fronto-striatal Functional Connectivity Supports Reward-Enhanced Memory in Older Adults"

**Whole-Brain Activation Analysis of MIE Task**

We conducted a whole-brain activation analysis during the anticipatory cue phase and separately during the target stimulus phase of the MIE task. In particular, we examined whether neural recruitment associated with subsequent memory would be modulated by reward magnitude and age. This analysis included only high confidence trials, with the 16 younger and 14 older adults described in the manuscript.

The first level models were created using the procedure described in the manuscript. Separate full factorial ANOVAs^[1]^ at the second level of analysis were used to examine activity during the reward anticipation and target stimulus phase of the experiment. Reward (high, low) and subsequent memory (hit, miss) were entered as within-subject factors and age as a between-subjects factor. Voxel threshold for all analyses was set at p < 0.005 (uncorrected), with a 15-voxel extent (k). MNI coordinates from SPM12 were converted to Talairach Coordinates using GingerAle (http://www.brainmap.org/ale). Anatomical labels in the cluster report tables were assigned using the Talairach Daemon (Lancaster et al., 1997) and visually checked using an anatomy atlas (Talairach and Tournoux, 1988).

**Results**

**Reward x Memory x Age Interaction at the Anticipation (Cue) Phase**

The fixed-effects mixed-ANOVA revealed a significant 3-way interaction in four regions (see Figure 1). This included a cluster in the superior frontal gyrus, posterior cingulate, inferior parietal lobule, and hippocampus. The activation was extracted from each of these clusters separately for each person. To uncover the pattern driving the three-way interaction in each region, percent signal change values from each cluster were submitted to separate mixed-ANOVAs with reward (high, low) and subsequent memory (hit [recognized], miss [forgotten]) as within-subjects variables and age (young, old) as a between-subjects variable. In each region identified, as expected there was a significant 3-way interaction, F(1,28) ≥ 7.71, p ≤ .01, η_p_^2^ ≥ .22. Follow-up 2 (Reward: high, low) x 2 (Memory: hit, miss) within-subjects ANOVAs were calculated for each age group separately. Results of these and follow-up tests are reported below for each region. See Table 1 for the full results of the ANOVA.

**Superior frontal gyrus**. The Reward x Memory interaction was significant for both younger, *F*(1, 15) = 7.46, *p* = .02, η_p_^2^ = .33, and older adults, *F*(1, 13) = 10.58, *p* = .05, η_p_^2^ = .22. Follow-up paired *t*-tests indicated that younger adults engaged the superior frontal gyrus to a greater extent during subsequently recognized (*M* = .50, *SE* = .13), compared to forgotten (*M* = .002, *SE* = .07), high-reward items *t*(15) = 3.14, *p* = .01, η^2^ = .40. During low-reward trials younger adults did not differentially engage the superior frontal gyrus for subsequently recognized (*M* = .10, *SE* = .13), compared to forgotten items (*M* = .52, *SE* = .08), *t*(15) = .37, *p* = .71, η^2^ = .009. Older adults showed the opposite pattern and engaged the superior frontal gyrus to a greater extent during subsequently recognized (*M* = .42, *SE* = .15), compared to forgotten (*M* = .12, *SE* = .12), low-reward items t(13) = 2.92, *p* = .01, η^2^ = .40, but did not differently engage this region for subsequently recognized (*M* = .13, *SE* = .12), compared to forgotten (*M* = .15, *SE* = .14), high-reward items, *t*(13) = .11, *p* = .91, η^2^ < .001.

**Posterior cingulate**. For younger adults the Reward x Memory interaction was significant, *F*(1, 15) = 25.58, *p* < .001, η_p_^2^ = .36, but for older adults the interaction was not significant, *F*(1, 13) = 1.51, *p* = .24. Follow-up paired *t*-tests with younger adults indicated there was a greater decrease in activity for subsequently recognized (*M* = -.23, *SE* = .24) compared to forgotten (*M* = -.04 *SE* = .10), high-reward items, *t*(15) = 3.14, *p* = .007, η^2^ = .40, but engaged the posterior cingulate to a greater extent for subsequently recognized (*M* = .12, *SE* = .15) compared to forgotten (*M* = -.04, *SE* = .19) low-reward items, *t*(15) = 3.86, *p* = .002, η^2^ = .50.

**Inferior parietal lobule**. For younger adults the Reward x Memory interaction was significant, *F*(1, 15) = 5.63, *p* = .03, η_p_^2^ = .24, but for older adults the interaction was not significant *F*(1, 13) = 3.59, *p* = .08. Follow-up paired *t*-tests revealed younger adults did not differently engage the inferior parietal lobule for subsequently recognized (*M* = .08, *SE* = .17), compared to forgotten items (*M* = -.20, *SE* = .11), high-reward items, *t*(15) = 1.40, *p* = .18, η^2^ = .12, but activity decreased in this region during subsequently recognized (*M* = -.39, *SE* = .18), compared to forgotten (*M* = .09, *SE* = .11), low-reward items, *t*(15) = 2.29, *p* = .04, η^2^ = .26.

**Hippocampus**. The Reward x Memory interaction was significant for both younger adults *F*(1, 15) = 10.99, *p* = .005, η_p_^2^ = .42, and older adults, *F*(1, 13) = 12.91, *p* = .003, η_p_^2^ = .50. Follow-up paired *t*-tests indicated that younger adults engaged the hippocampus to a greater extent during subsequently recognized (*M* = .17, *SE* = .17), compared to subsequently forgotten (*M* = -.42, *SE* = .16), high-reward items, *t*(15) = 3.05, *p* = .008, η^2^ = .38, but did not differentially engage this region for subsequently recognized (*M* = -.16, *SE* = .15), compared to forgotten (*M* = .06, *SE* = .13), low-reward items, *t*(15) = 1.53, *p* = .15, η^2^ = .13. Follow-up *t*-tests with older adults revealed the opposite pattern. Older adults did not differentially engage the hippocampus during subsequently recognized (*M* = -.21, *SE* = .16) compared to subsequently forgotten (*M* = .05, *SE* = .16) high-reward items, *t*(13) = 1.59, *p* = .14, η^2^ = .16, but engaged it to a greater extent for subsequently recognized (*M* = .24, *SE* = .20) compared to forgotten (*M* = -.20, *SE* = .13), low-reward items, *t*(13) = 2.42, *p* = .03, η^2^ = .31.

**Reward x Memory x Age interaction at the Target Stimulus Phase**

Next, we were interested in examining the activation patterns during the target stimulus phase. In particular, whether neural recruitment associated with subsequent memory would be modulated by the reward value and age group. The fixed-effects mixed-ANOVA revealed a significant 3-way interaction in four regions: superior frontal gyrus, cingulate gyrus and bilateral insula (see Figure 2). The activation was extracted from each of these clusters separately for each person. To uncover the pattern driving the three-way interaction in each region, percent signal change values from each cluster were submitted to separate mixed-ANOVAs with reward (high, low) and subsequent memory (hit, miss) as within-subjects variables and age (young, old) as a between-subjects variable. In each region identified, as expected there was a significant 3-way interaction, *F*(1,28) ≥ 16.51, *p* ≤ .001, η_p_^2^ ≥ .37. A Follow-up 2 (Reward: high, low) x 2 (Memory: hit, miss) within-subjects ANOVAs were calculated for each age group separately. Results of these follow-up tests are reported below for each region. See Table 2 for the full results of the ANOVA.

**Superior frontal gyrus**. The Reward x Memory interaction was not significant for younger adults, *F*(1, 15) = 3.51, *p* = .08, but was significant for older adults, *F*(13) = 12.23, *p* = .004, η_p_^2^ = .49. Follow-up paired *t*-tests revealed that older adults did not differently engage this region for subsequently recognized *(M* = -.14, *SE* = .11) compared to forgotten (*M* = .09, *SE* = .12) high-reward items, *t*(13) = 2.03, *p* = .06, η^2^ = .24, but engaged this region to a greater extent for subsequently recognized (*M* = .35, *SE* = .22) compared to forgotten (*M* = -.20, *SE* = .11) low-reward items, *t*(13) = 3.24, *p* = .006, η^2^ = .44.

**Cingulate gyrus***.* The Reward x Memory interaction was significant for both younger, *F*(1, 15) = 5.35, *p* = .04, η_p_^2^ = .26, and older adults, *F*(1, 13) = 5.72, *p* = .03, η_p_^2^ = .31. Follow-up paired *t*-tests indicated that younger adults did not differently engaged this region for subsequently recognized (*M* = .32, *SE* = .12), compared to forgotten (*M* = .01, *SE* = .13), high-reward items *t*(15) = 1.39, *p* = .19, η^2^ = .11, nor during subsequently recognized (*M* = .02, *SE* = .14), compared to forgotten (*M* = .29, *SE* = .13), low-reward items *t*(15) = 1.53, *p* = .15, η^2^ = .13. Older adults exhibited a decrease in activity during subsequently recognized (*M* = -.47, *SE* = .18) compared to forgotten (*M* = .09, *SE* = .09) high-reward items, *t*(13) = 2.34, *p* = .04, η^2^ = .03, but there was no difference in activation for subsequently recognized (*M* = .16, *SE* = .16) compared to forgotten (*M* = .30, *SE* = .03) low-reward items, *t*(13) = 1.24, *p* = .24, η^2^ = .11.

**Left insula**. The Reward x Memory interaction was significant for both younger, *F*(1, 15) = 14.39, *p* = .002, η_p_^2^ = .49, and older adults, *F*(1, 13) = 7.52, *p* = .02, η_p_^2^ = .31. Follow-up paired samples *t*-tests indicated that younger adults engaged this region to a greater extent during subsequently recognized (*M* = .25, *SE* = .13) compared to forgotten (*M* = -.26, *SE* = .10), high-reward items, *t*(15) = 3.36, *p* = .004, η^2^ = .43, and subsequently recognized (*M* = .32, *SE* = .13) compared to forgotten (*M* = .02, *SE* = .17), low-reward items, *t*(15) = 2.24, *p* = .04, η^2^ = .25. Older adults showed less activation in this region during subsequently recognized (*M* = -.34, *SE* = .10) compared to forgotten (*M* = .16, *SE* = .11), high-reward items, *t*(13) = 3.71, *p* = .04, η^2^ = .51, but no differences in activation for subsequently recognized (*M* = -.09, *SE* = .11) compared to forgotten (*M* = -.19, *SE* = .11), low-reward items, *t*(13) = .63, *p* = .54, η^2^ = .03.

**Right insula**. The right insula showed a slightly different pattern than the left insula. For younger adults the Reward x Memory interaction was significant, *F*(15) = 15.55, *p* < .001, η_p_^2^ = .51, but for older adults the interaction was not significant, *F*(1, 13) = 2.96, *p* = .11, η_p_^2^ = .51. Follow-up paired *t*-tests with younger adults revealed they recruited this region to a greater extent for subsequently recognized (*M* = .46, *SE* = .20) compared to forgotten (*M* = -.20, *SE* = .09) high-reward items, *t*(15) = 3.31, *p* = .005, η^2^ = .42. The *t-*test for low-reward trials was marginally significant, *t*(15) = 2.04, *p* = .06, η^2^ = .22, and indicated that right insula was recruited less for subsequently recognized (*M* = -.21, *SE* = .19) compared to forgotten items (*M* = .21, *SE* = .13).

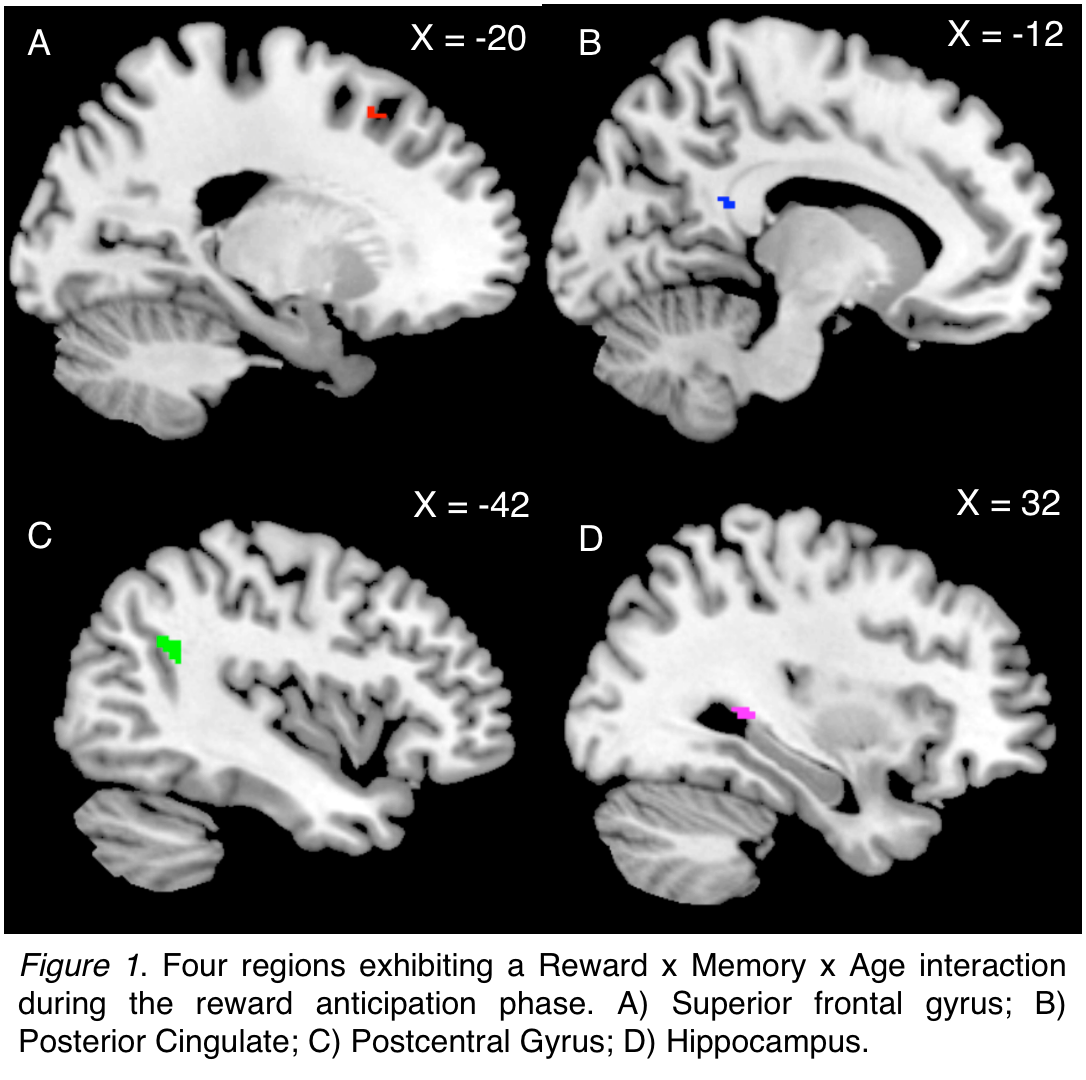

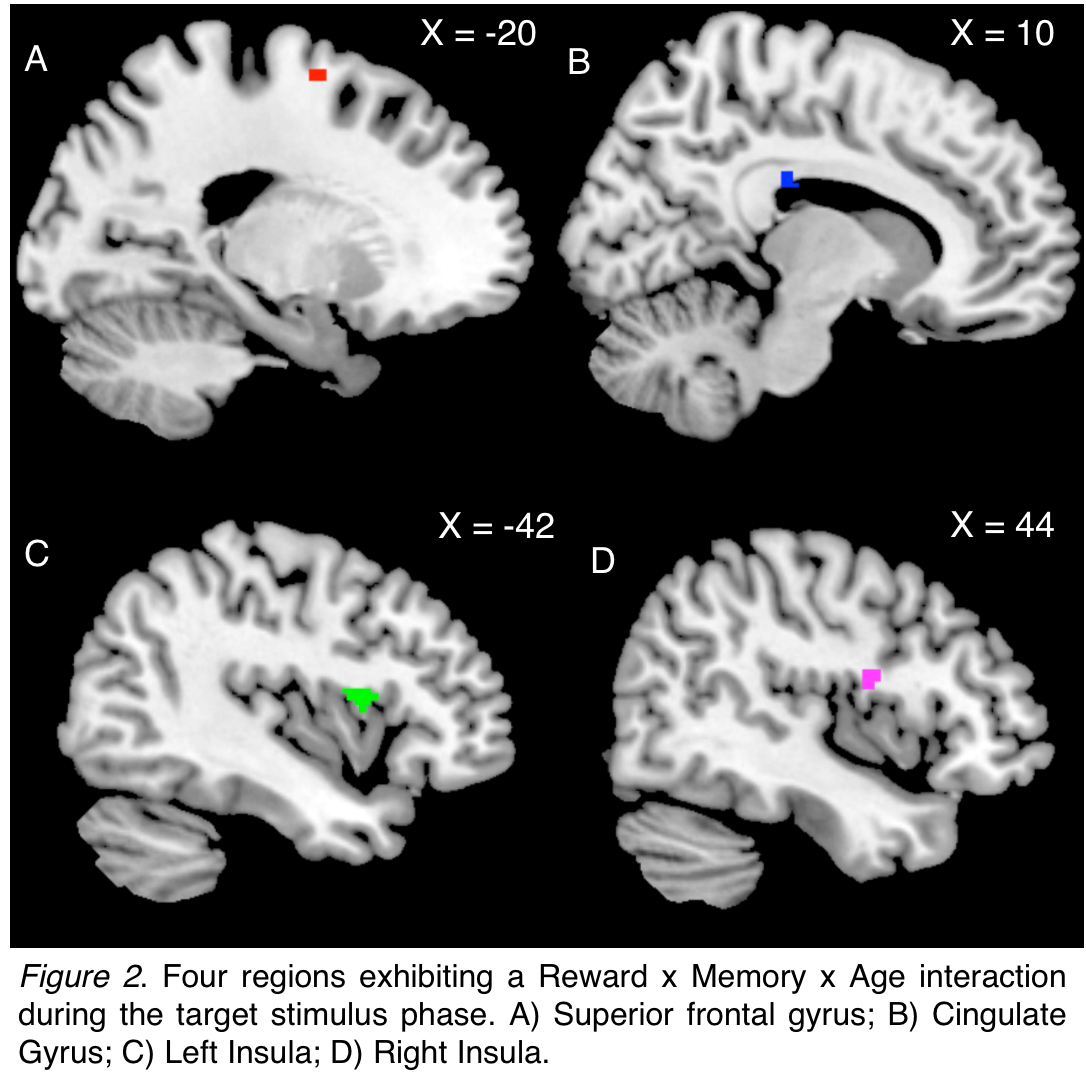

Table 1

*Regions from the whole-brain activation analysis during anticipation phase of the Monetary Incentive Encoding Task*

| Lobe | | Region | Hem | BA | MNI Coordinates  X Y Z | T-value | k |
| --- | --- | --- | --- | --- | --- | --- | --- |
| *Main Effect of Memory* | | | | | | | |
| Cerebellum |  | | R | * | 22 -32 -24 | 5.23 | 724 |
| Frontal | Inferior Frontal Gyrus | | L | 47 | -38 34 -12 | 5.2 | 111 |
|  |  | |  | 9 | -36 12 26 | 4.08 | 70 |
|  | Middle Frontal Gyrus | | L | 46 | -48 26 18 | 3.66 | 86 |
|  | Precentral Gyrus | | R | 6 | 58 4 38 | 4.09 | 36 |
|  | Superior Frontal Gyrus | | L | 8 | -18 36 44 | 4.34 | 199 |
| Limbic | Cingulate Gyrus | | L | 24 | -16 -12 48 | 4.59 | 21 |
|  | Amygdala | | R | * | 32 -4 -16 | 5.77 | 51 |
|  | Posterior Cingulate | | L | 30 | -8 -54 12 | 4.08 | 63 |
|  |  | | R | 30 | 16 -52 12 | 3.56 | 25 |
| Parietal | Precuneus | | L | 31 | -24 -72 32 | 3.36 | 22 |
| Other | Caudate Body | | L | * | -12 2 28 | 3.47 | 29 |
|  | Caudate Tail | | R | * | 32 -44 12 | 4.37 | 16 |
|  | Putamen | | R | * | 36 -20 -12 | 3.43 | 64 |
|  | Thalamus | | L | * | 0 2 2 | 3.51 | 23 |
| Temporal | Fusiform Gyrus | | L | 37 | -46 -66 -12 | 5.56 | 832 |
|  | Middle Temporal Gyrus | | L | 39 | -40 -78 22 | 4.93 | 439 |
|  | No Gray Matter | |  | * | 38 -68 24 | 4.6 | 453 |
| *Main Effect of Reward* | | | | | | | |
| Cerebellum |  | | L | * | -26 -52 -14 | 4.22 | 323 |
|  |  | |  | * | -12 -50 -8 | 3.39 | 27 |
|  |  | | R | * | 20 -32 -22 | 3.61 | 39 |
|  |  | |  | * | 24 -56 -10 | 3.35 | 40 |
| Frontal | Inferior Frontal Gyrus | | R | 9 | 42 10 26 | 3.51 | 124 |
|  | Medial Frontal Gyrus | | R | 6 | 10 10 52 | 3.53 | 67 |
|  | Precentral Gyrus | | L |  | -46 -2 30 | 4.96 | 867 |
|  |  | |  |  | -36 -6 44 | 4.86 | 56 |
|  |  | | R | 4 | 36 -14 66 | 3.78 | 234 |
|  | No gray matter | | L | 6 | -20 0 58 | 3.86 | 52 |
| Limbic | Cingulate Gyrus | | L | 32 | -4 26 28 | 3.32 | 29 |
|  |  | |  | 24 | -6 12 44 | 3.22 | 33 |
|  |  | | R | 24 | 4 12 28 | 3.71 | 17 |
|  | Parahippocampal Gyrus | | R | 36 | 26 -42 -6 | 3.24 | 25 |
|  | Posterior Cingulate | | L | 30 | -2 -54 8 | 3.39 | 20 |
| Occipital | Fusiform Gyrus | | R | 19 | 34 -84 -10 | 3.12 | 34 |
|  | Middle Occipital Gyrus | | L |  | -34 -86 18 | 3.38 | 65 |
|  |  | | R |  | 38 -82 14 | 3.61 | 140 |
|  |  | |  |  | 46 -68 10 | 3.18 | 18 |
| Parietal | Inferior Parietal Lobule | | L | 40 | -32 -36 42 | 3.49 | 20 |
|  | Postcentral Gyrus | | L | 2 | -56 -20 34 | 4.56 | 138 |
|  |  | | R |  | 52 -18 46 | 3.8 | 31 |
|  |  | |  |  | 56 -22 32 | 3.35 | 18 |
|  | Precuneus | | L | 7 | -22 -62 38 | 3.65 | 58 |
|  |  | | R |  | 20 -56 56 | 2.9 | 19 |
|  | Superior Parietal Lobule | | L | 7 | -30 -46 46 | 3.48 | 30 |
|  | Superior Parietal Lobule | | L | 7 | -28 -56 48 | 2.98 | 25 |
|  | Caudate Body | | L | * | -14 12 14 | 3.23 | 95 |
|  |  | | R | * | 12 8 6 | 3.35 | 75 |
| Other | Claustrum | | L | * | -30 10 4 | 3.83 | 62 |
|  | Putamen | | L | * | -26 -2 20 | 3.22 | 20 |
|  |  | | R | * | 14 0 -12 | 3.33 | 19 |
|  |  | |  | * | 32 14 0 | 3.05 | 18 |
|  | Thalamus | | L | * | 0 -4 2 | 3.66 | 62 |
|  |  | |  | * | -8 -12 6 | 3.27 | 31 |
|  |  | | R | * | 14 -26 0 | 3.41 | 28 |
|  |  | |  | * | 16 -12 8 | 3.28 | 21 |
| Temporal | Fusiform Gyrus | | L | 37 | -48 -58 -8 | 4.16 | 76 |
|  |  | | R |  | 36 -48 -14 | 3.86 | 26 |
|  |  | |  |  | 42 -58 -10 | 3.52 | 21 |
|  | Middle Temporal Gyrus | | L | 39 | -42 -78 24 | 3.08 | 20 |
| *Main Effect of Age* | | | | | | | |
| Occipital | Cuneus | | R | 17 | 8 -84 12 | 3.94 | 30 |
| *Memory x Reward Interaction* | | | | | | | |
| Cerebellum |  | | L | * | -6 -30 -28 | 3.77 | 41 |
|  |  | |  | * | -14 -44 -22 | 3.41 | 21 |
| Frontal | Middle Frontal Gyrus | | L | 9 | -34 24 32 | 3.79 | 90 |
|  | No gray matter | | L | 6 | -18 0 60 | 4.32 | 90 |
| Limbic | Cingulate Gyrus | | L | 32 | -14 16 42 | 3.88 | 48 |
|  |  | |  | 31 | -18 -20 36 | 3.25 | 19 |
|  |  | | R | 24 | 12 8 38 | 3 | 20 |
|  | Parahippocampal Gyrus | | R | 30 | 18 -36 -6 | 3.85 | 93 |
|  | Posterior Cingulate | | L | 29 | -2 -40 16 | 3.45 | 22 |
| Occipital | Lingual Gyrus | | L | * | -20 -78 4 | 3.72 | 26 |
| Parietal | Precuneus | | L | 7 | -24 -56 56 | 3.46 | 52 |
|  |  | |  |  | -16 -46 58 | 3.32 |  |
|  |  | | R |  | 26 -58 56 | 3.6 | 27 |
| Other | Caudate Body | | L | * | -14 14 10 | 3.09 | 23 |
|  |  | | R | * | 12 16 6 | 3.64 | 17 |
|  | Claustrum | |  | * | 42 -2 -4 | 3.19 | 16 |
|  | Insula | | L | 13 | -32 16 8 | 3.06 | 16 |
|  |  | | R |  | 34 -30 14 | 3.91 | 49 |
|  | Putamen | | L | * | -24 4 -14 | 3.57 | 28 |
|  | Thalamus | | L | * | -30 -32 4 | 4.43 | 19 |
|  |  | |  | * | -24 -30 14 | 3.96 | 23 |
|  |  | | R | * | 16 -28 2 | 3.7 |  |
| Temporal | Caudate Tail | | R | * | 42 -30 2 | 4.22 | 28 |
|  |  | |  | * | 40 -38 2 | 3.85 | 18 |
|  | No gray matter | | L | 21 | -42 -6 -12 | 3.32 | 15 |
| *Memory x Age Interaction* | | | | | | | |
| No significant Clusters | | | | | | | |
| *Age x Reward Interaction* | | | | | | | |
| Cerebellum |  | | L | * | -42 -48 -44 | 3.74 | 22 |
|  |  | | L | * | -30 -68 -48 | 3.44 | 24 |
|  |  | | R | * | 24 -80 -26 | 3.1 | 17 |
| Frontal | Medial Frontal Gyrus | | L | 9 | -12 36 30 | 3.53 | 18 |
|  |  | | R | 6 | 14 38 32 | 3.58 | 40 |
|  |  | |  | 8 | 4 50 34 | 3.44 | 47 |
| Limbic | Anterior Cingulate | | R | 32 | 20 44 6 | 4.31 | 59 |
|  | Cingulate Gyrus | | L |  | -16 20 24 | 4.63 | 19 |
|  | Parahippocampal Gyrus | | L | 28 | -18 -26 -24 | 3.61 | 17 |
|  |  | | R | 19 | 32 -48 -8 | 3.83 | 24 |
| Other | Caudate Tail | | R | * | 26 -34 18 | 3.86 | 18 |
| *Reward x Memory x Age Interaction* | | | | | | | |
| Frontal | Superior Frontal Gyrus | | L | 6 | -20 20 50 | 3.16 | 18 |
| Limbic | Posterior Cingulate | | L | 29 | -12 -44 22 | 3.16 | 16 |
| Parietal | Inferior Parietal Lobule | | L | 40 | -42 -50 26 | 3.07 | 25 |
| Temporal | Hippocampus | | R | * | 32 -38 4 | 3.56 | 16 |

Note. Hem = hemisphere; L = Left; R = Right; BA = Brodmann Area; * = no Brodmann Area; MNI = Montreal Neurological Institute; k = cluster extent.

Table 2

*Regions from the whole-brain activation analysis during target stimulus phase of the Monetary Incentive Encoding Task*

| Lobe | Region | Hem | BA | MNI Coordinates  X Y Z | T-value | k |
| --- | --- | --- | --- | --- | --- | --- |
| *Main Effect of Memory* | | | | | | |
| Frontal | Precentral Gyrus | L | 6 | -56 2 30 | 3.52 | 31 |
| Limbic | Cingulate Gyrus | L | 24 | -14 -6 34 | 3.44 | 29 |
| Parietal | Postcentral Gyrus | R | 2 | 54 -18 32 | 3.68 | 41 |
| Other | Claustrum | L | * | -36 -4 -6 | 3.58 | 17 |
| *Main Effect of Reward* | | | | | | |
| Cerebellum |  | L | * | -16 -44 -28 | 3.92 | 25 |
|  |  |  | * | -10 -48 2 | 4.07 | 74 |
|  |  |  | * | -42 -54 -28 | 3.31 | 18 |
|  |  |  | * | -20 -40 -44 | 4.26 | 52 |
|  |  |  | * | -44 -50 -42 | 3.77 | 114 |
|  |  |  | * | -2 -60 -40 | 3.36 | 15 |
|  |  |  | * | -26 -76 -46 | 3.36 | 25 |
|  |  | R | * | 10 -32 -22 | 3.84 | 94 |
|  |  |  | * | 28 -38 -24 | 3.46 | 27 |
|  |  |  | * | 34 -68 -34 | 4.22 | 509 |
|  |  |  | * | 24 -36 -36 | 3.13 | 16 |
|  |  |  | * | 32 -84 -14 | 4.58 | 206 |
|  |  |  | * | 10 -78 -46 | 3.29 | 36 |
|  |  |  | * | 16 -82 -42 | 3 |  |
|  |  |  | * | 12 -84 -28 | 4.28 | 67 |
| Frontal | Cingulate Gyrus | R | 32 | 16 30 34 | 4.01 | 41 |
|  |  |  |  | 18 18 32 | 3.26 | 28 |
|  | Inferior Frontal Gyrus | L | * | -42 50 -10 | 4.87 | 394 |
|  |  |  | 9 | -50 12 32 | 3.51 | 28 |
|  |  | R | 46 | 48 44 -6 | 3.65 | 32 |
|  |  |  | 45 | 38 28 -2 | 3.14 | 20 |
|  | Medial Frontal Gyrus | L | 10 | -20 58 -2 | 3.59 | 35 |
|  |  |  | 32 | -24 44 0 | 3.44 | 19 |
|  |  | R | 6 | 16 -12 60 | 3.51 | 17 |
|  |  |  | 9 | 10 48 14 | 3.41 | 51 |
|  | Middle Frontal Gyrus | L | 46 | -48 30 24 | 4.17 | 34 |
|  |  |  | 9 | -32 26 28 | 4.1 | 61 |
|  |  |  | 6 | -30 12 56 | 3.44 | 173 |
|  |  | R | 9 | 44 28 28 | 4.59 | 364 |
|  |  |  |  | 56 12 34 | 3.93 | 128 |
|  |  |  | 6 | 30 12 40 | 3.87 | 68 |
|  |  |  |  | 34 16 56 | 3.22 | 16 |
|  |  |  | 11 | 32 40 -14 | 3.81 | 67 |
|  | Precentral Gyrus | L | 9 | -42 26 36 | 3.93 | 101 |
|  |  |  | 3 | -56 -14 26 | 3.44 | 36 |
|  |  |  | 6 | -42 2 24 | 3.19 | 72 |
|  | Subcallosal Gyrus | R | 34 | 28 4 -16 | 4 | 21 |
|  | Superior Frontal Gyrus | L | 8 | -20 38 48 | 3.28 | 86 |
|  |  |  |  | 2 40 46 | 3.1 | 31 |
| Limbic | Anterior Cingulate | L | 32 | -16 38 14 | 3.75 | 29 |
|  |  | R | 24 | 14 26 14 | 3.95 | 64 |
|  | Cingulate Gyrus | L | 32 | -20 16 36 | 3.55 | 53 |
|  |  |  | 24 | -16 0 40 | 3.35 | 18 |
|  |  | R | 31 | 8 -34 36 | 3.41 | 16 |
|  | Parahippocampal Gyrus | L | 36 | -44 -24 -18 | 4.43 | 80 |
|  |  |  |  | -26 -30 -18 | 3.41 | 29 |
|  |  | R |  | 50 -34 -12 | 3.73 | 23 |
|  | Posterior Cingulate | L | 31 | -28 -70 24 | 3.72 | 90 |
|  |  | R | 30 | 22 -56 8 | 4.11 | 445 |
| Occipital | Cuneus |  | 19 | 30 -86 28 | 3.88 | 272 |
|  | Lingual Gyrus | L | 17 | -12 -100 -4 | 3.96 | 40 |
|  |  |  | 18 | -12 -68 0 | 3.59 | 23 |
|  |  |  | 19 | -30 -80 -4 | 3.55 | 48 |
|  |  | R |  | 34 -62 6 | 3.74 | 26 |
|  | Middle Occipital Gyrus | L | 19 | -32 -90 20 | 3.56 | 172 |
| Parietal | Inferior Parietal Lobule | L | 40 | -32 -28 40 | 4.09 | 24 |
|  |  |  |  | -46 -56 46 | 3.27 | 24 |
|  | Postcentral Gyrus | R | 3 | 56 -18 40 | 3.69 | 18 |
|  | Precuneus | L | 7 | -20 -64 32 | 3.5 | 71 |
|  |  | R | 31 | 30 -72 34 | 3.64 | 123 |
|  | Superior Parietal Lobule | L | 7 | -28 -60 46 | 3.52 | 61 |
|  |  | R |  | 32 -58 50 | 3.09 | 21 |
|  | Supramarginal Gyrus | L | 40 | -40 -40 40 | 3.87 | 77 |
| Other | Caudate Body | L | * | -16 14 4 | 3.82 | 40 |
|  |  |  | * | -22 -2 28 | 3.34 | 15 |
|  | Caudate Tail | L | * | -22 -30 20 | 3.12 | 15 |
|  | Caudate Head | R | * | 10 20 -2 | 3.72 | 20 |
|  | Caudate Body | R | * | 12 4 14 | 3.45 | 17 |
|  | Claustrum | L | * | -28 -4 16 | 3.27 | 18 |
|  | Insula | R | * | 50 -22 18 | 3.55 | 22 |
|  | Putamen | R | * | 26 8 0 | 3.59 | 17 |
| Temporal | Fusiform Gyrus | L | 37 | -30 -42 -16 | 3.82 | 229 |
|  | Middle Temporal Gyrus | L | 37 | -46 -62 0 | 4.26 | 451 |
|  | Superior Temporal Gyrus | R | 41 | 44 -30 8 | 4.24 | 50 |
|  |  | R |  | 60 -30 8 | 3.45 | 21 |
| *Main Effect of Age* | | | | | | |
| Cerebellum |  | L | * | -18 -42 -20 | 3.73 | 45 |
|  |  |  | * | -42 -74 -40 | 3.02 | 17 |
|  |  |  | * | 0 -82 -30 | 3.11 | 15 |
|  |  | R | * | 22 -78 -12 | 4.3 | 422 |
| Frontal | Inferior Frontal Gyrus | R | 47 | 42 38 -16 | 3.02 | 19 |
|  | Superior Frontal Gyrus | L | 9 | -10 62 20 | 3.64 | 54 |
|  |  |  | 8 | -20 32 44 | 3.22 | 29 |
|  |  | R |  | 24 50 36 | 3.29 | 36 |
| Limbic | Anterior Cingulate | R | 32 | 8 50 -8 | 3.86 | 296 |
|  | Cingulate Gyrus | L | 31 | -8 -44 32 | 3.14 | 28 |
|  | Parahippocampal Gyrus | L | 30 | -14 -38 6 | 3.82 | 54 |
|  | Posterior Cingulate | L | 31 | -30 -64 24 | 3.88 | 96 |
|  |  |  | 30 | -16 -56 10 | 3.44 | 34 |
|  |  | R | 23 | 6 -54 18 | 3.42 | 90 |
| Occipital | Fusiform Gyrus | L | 19 | -38 -84 -8 | 3.25 | 16 |
|  | Lingual Gyrus | L | 17 | -16 -92 10 | 4.73 | 780 |
|  | Middle Occipital Gyrus | L | 19 | -36 -82 14 | 4.05 | 94 |
|  |  | R |  | 36 -82 18 | 3.79 | 249 |
| Parietal | Precuneus | L | 7 | -10 -72 46 | 3.12 | 19 |
| Other | Caudate Body | L | * | -20 24 16 | 3.63 | 39 |
|  | Claustrum | R | * | 40 -6 -8 | 3.22 | 17 |
|  | Insula | R | 13 | 32 30 12 | 3.3 | 15 |
| Temporal | Superior Temporal Gyrus | R | 38 | 38 0 -28 | 3.5 | 32 |
|  | No gray matter |  | *** | 20 2 30 | 3.48 | 22 |
| *Memory x Reward Interaction* | | | | | | |
| Cerebellum |  | L | * | -20 -50 -40 | 3.36 | 30 |
|  |  | R | * | 36 -68 -32 | 4.27 | 43 |
| Frontal | Inferior Frontal Gyrus | R | 45 | 54 26 10 | 3.57 | 15 |
| Limbic | Anterior Cingulate | L | 32 | -16 38 14 | 3.76 | 30 |
| Limbic | Cingulate Gyrus | L | 31 | -22 -32 48 | 4.33 | 86 |
|  | Parahippocampal Gyrus | L | 36 | -42 -22 -18 | 3.96 | 65 |
|  |  |  |  | -28 -30 -18 | 3.78 | 29 |
| Occipital | Middle Temporal Gyrus | L | 19 | -34 -62 20 | 3.16 | 22 |
| Parietal | Precuneus | R | 7 | 18 -44 52 | 3.56 | 19 |
| Other | Claustrum | L | * | -38 0 -12 | 3.69 | 22 |
|  | Insula | L | 13 | -30 -32 14 | 3.61 | 48 |
|  | Thalamus | R | * | 8 -36 10 | 4.05 | 28 |
| Temporal | Caudate Tail | R | * | 42 -18 -18 | 4.26 | 52 |
|  | Fusiform Gyrus | L | 37 | -42 -46 -12 | 4.7 | 47 |
|  |  |  |  | -38 -60 -8 | 3.9 | 136 |
|  | No gray matter | L | 37 | -52 -50 -10 | 3.15 | 26 |
|  |  | R | 21 | 46 -6 -20 | 3.38 | 39 |
| *Memory x Age Interaction* | | | | | | |
| Cerebellum |  | L | * | 0 -44 -38 | 3.52 | 31 |
|  |  |  | * | -58 -44 40 | 3.6 | 38 |
|  |  | R | * | 14 -58 -26 | 3.42 | 19 |
|  |  |  | * | 10 -60 30 | 2.86 | 18 |
| Frontal | Inferior Frontal Gyrus | L | 45 | -56 20 20 | 4.52 | 48 |
|  | Middle Frontal Gyrus | L | 6 | -28 4 52 | 3.63 | 23 |
|  |  |  |  | -18 -12 52 | 3.45 | 23 |
|  |  |  |  | -14 4 62 | 3.36 | 15 |
|  | Middle Frontal Gyrus | R | 9 | 40 36 32 | 3.13 | 17 |
|  | Precentral Gyrus | L | 6 | -58 2 12 | 4.22 | 60 |
|  |  |  |  | -44 0 54 | 3.45 | 21 |
|  |  | R | 4 | 64 -12 34 | 3.71 | 25 |
|  | Superior Frontal Gyrus | L | 6 | -8 20 50 | 3.9 | 41 |
|  |  |  |  | -16 18 60 | 3.82 | 24 |
|  |  |  | 9 | -20 42 30 | 3.88 | 65 |
|  |  |  |  | -8 60 16 | 3.4 | 26 |
|  |  | R |  | 18 54 26 | 3.2 | 22 |
| Limbic | Anterior Cingulate | L | 32 | -10 48 -6 | 3.93 | 253 |
|  |  | L | 24 | -10 36 12 | 3.43 | 66 |
|  |  | R | * | 4 34 -8 | 3.65 | 20 |
|  | Cingulate Gyrus | L | 31 | -4 -38 42 | 3.94 | 41 |
|  |  |  | 32 | -12 26 42 | 3.35 | 47 |
|  |  |  |  | -6 16 42 | 3.15 | 16 |
|  |  | R | 24 | 14 10 30 | 3.59 | 18 |
|  | Parahippocampal Gyrus | R | * | 36 -6 -26 | 3.38 | 21 |
|  | Uncus | L | * | -30 0 -28 | 3.66 | 15 |
| Occipital | Cuneus | L | 17 | -16 -82 14 | 3.24 | 28 |
|  | Lingual Gyrus | R | * | 20 -74 2 | 3.55 | 40 |
|  | Precuneus | L | 31 | -4 -66 34 | 4.3 | 481 |
| Other | Claustrum | L | * | -24 24 -10 | 3.84 | 25 |
|  |  |  | * | -34 -6 6 | 3.29 | 15 |
|  | Insula | L | 13 | -40 10 -8 | 3.61 | 22 |
|  | Putamen | L | * | -30 4 6 | 4.31 | 49 |
|  |  |  | * | -20 2 12 | 4.21 | 17 |
|  | Thalamus | R | * | 6 -10 -4 | 3.25 | 18 |
| Temporal | Angular Gyrus | L | 39 | -36 -76 38 | 3.18 | 18 |
|  | Superior Temporal Gyrus | L | 39 | -42 -58 36 | 3.17 | 38 |
|  |  | R | 22 | 56 -2 -6 | 3.22 | 15 |
| *Age x Reward Interaction* | | | | | | |
| Parietal | Precuneus | R | 31 | 22 -40 34 | 3.8 | 18 |
|  |  | R | 19 | 34 -60 42 | 3.63 | 51 |
| Other | Putamen | R | * | 34 -20 -6 | 3.56 | 15 |
|  | Thalamus | R | * | 26 -20 18 | 3.95 | 39 |
| *Reward x Memory x Age Interaction* | | | | | | |
| Frontal | Superior Frontal Gyrus | L | 6 | -20 -2 64 | 3.78 | 18 |
| Limbic | Cingulate Gyrus | R | 23 | 10 -24 26 | 3.83 | 19 |
| Other | Insula | L | 13 | -42 10 10 | 3.61 | 46 |
|  | Insula | R |  | 44 0 18 | 3.59 | 15 |

Note. Hem = hemisphere; L = Left; R = Right; BA = Brodmann Area; * = no Brodmann Area; MNI = Montreal Neurological Institute; k = cluster extent.

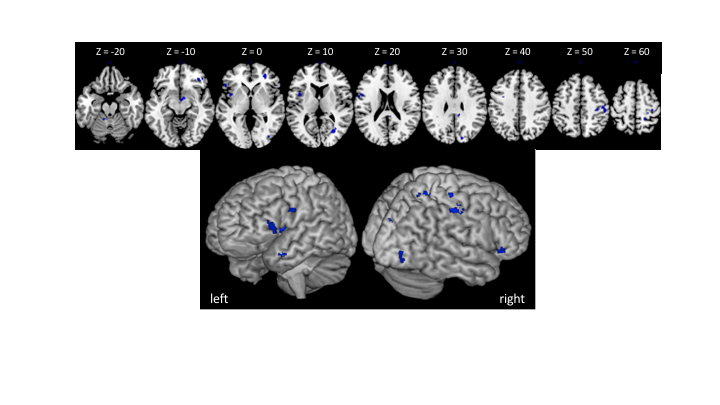

*Figure 3.* All Regions exhibiting a Reward x Memory x Age interaction in connectivity with the caudate. Axial slices labelled with MNI Z-coordinate (top panel). Whole brain render from left and right hemisphere perspective (bottom panel).

1. Results of this whole-brain activation analysis are reported in the supplemental material.
